# Lamprey Lecticans Link New Vertebrate Genes to the Origin and Elaboration of Vertebrate Tissues

**DOI:** 10.1101/2020.09.24.311837

**Authors:** Zachary D. Root, David Jandzik, Cara Allen, Margaux Brewer, Marek Romášek, Tyler Square, Daniel M. Medeiros

## Abstract

The evolution of vertebrates from an invertebrate chordate ancestor involved the evolution of new organs, tissues, and cell types. It was also marked by the origin and duplication of new gene families. If, and how, these morphological and genetic innovations are related is an unresolved question in vertebrate evolution. Hyaluronan is an extracellular matrix (ECM) polysaccharide important for water homeostasis and tissue structure. Vertebrates possess a novel family of hyaluronan binding proteins called Lecticans, and studies in jawed vertebrates (gnathostomes) have shown they function in many of the cells and tissues that are unique to vertebrates. This raises the possibility that the origin and/or expansion of this gene family helped drive the evolution of these vertebrate novelties. In order to better understand the evolution of the *lectican* gene family, and its role in the evolution of vertebrate morphological novelties, we investigated the phylogeny, genomic arrangement, and expression patterns of all *lecticans* in the sea lamprey (*Petromyzon marinus*), a jawless vertebrate. Though both *P. marinus* and gnathostomes have four *lecticans*, our phylogenetic and syntenic analyses suggest lamprey *lecticans* are the result of one or more cyclostome-specific duplications. Despite the independent expansion of the lamprey and gnathostome *lectican* families, we find highly conserved expression of *lecticans* in vertebrate-specific and mesenchyme-derived tissues. We also find that, unlike gnathostomes, lamprey expresses its *lectican* paralogs in distinct subpopulations of head skeleton precursors, potentially reflecting an ancestral diversity of skeletal tissue types. Together, these observations suggest that the ancestral pre-duplication *lectican* had a complex expression pattern, functioned to support mesenchymal histology, and likely played a role in the evolution of vertebrate-specific cell and tissue types.

## INTRODUCTION

The emergence of vertebrates involved the elaboration of the ancestral chordate body plan with an array of new cell types, tissues, and organs. Among these are the expanded central and peripheral nervous systems, and the complex skeletomuscular systems of the head and trunk, which includes an array of new structural and connective tissues [1, 2]. Interestingly, large portions of these novelties are derived from the same embryonic source, neural crest cells, which also give rise to parts of the heart, teeth, endocrine system, and vascular smooth muscle [1–3]. The evolution of these morphological and developmental novelties coincided with major genome-wide changes including the origin of several new gene families, at least one whole genome duplication, and the evolution of new gene regulatory networks [4–15]. The timing of these genomic events has led to speculation that they facilitated the origin and morphological diversification of vertebrates by altering early development.

While alterations in embryogenesis can lead to major changes in the body plan, the evolution of truly novel tissues and cell types also requires the evolution of new cellular functions and histological properties. Extracellular matrix (ECM) proteins not only provide support and structure to cells and tissues, but also mediate signal transduction and mechanotransduction [16]. A key component of the ECM of many vertebrate tissues is a vertebrate-specific family of proteoglycans called Lecticans. Structurally, Lecticans are complex, consisting of hyaluronan-binding X-link domains, c-type lectin domains, a chondroitin/keratan sulfate binding domain, and an immunoglobulin domain. Because of this modular structure, Lecticans are able to interface with many different types of molecules and perform a range of functions in the ECM of diverse cells and tissues [17].

Genomically, all gnathostome *lectican* paralogs are closely linked to a *hapln* gene, which also encodes an X-link domain containing protein [18]. The proximity of *lecticans* and *haplns*, together with their high sequence similarity, indicate they evolved via tandem duplication of an ancestral X-link protein-encoding gene, with subsequent exon shuffling resulting in the hybrid structure of Lecticans [19]. After assembly of the primordial *lectican* gene, two genome-wide duplications are thought to have generated the four paralogs seen in modern jawed vertebrates: *aggrecan* (*acan*), *brevican* (*bcan*), *neurocan* (*ncan*), and *versican* (*vcan*). Since these duplications, the structures of the four gnathostome Lecticans have diverged, with *acan* acquiring an additional, X-link domain, *bcan* and *ncan* losing an interglobular fold sequence adjacent to the immunoglobulin domain, and all Lecticans evolving chondroitin/keratan sulfate binding domains of different sizes [17].

Subfunctionalization, specialization, and/or neofunctionalization of gnathostome *lectican* paralogs resulted in each possessing distinct expression patterns and functions, in neural, skeletal, cardiac, and connective tissues [20, 21]. *acan* is known primarily for its role in the cartilage ECM [22, 23] [among others], but it is also involved in neural crest cell migration and synaptic complexes in the brain [24–28]. *acan* expression has also been found in the developing notochord as well as the epicardium and mesenchyme of the heart [29, 30]. *vcan* is the most widely expressed *lectican* and is transcribed in mesoderm-derived tissues and organs including the kidneys, heart, muscles, and skeleton [29–34], and various neurectodermal derivatives like the otic vesicle, lens primordium [32], oligodendrocytes, Schwann Cells, the perineuronal net, ectodermal placodes, and migrating neural crest cells [20, 28, 35–38]. *bcan* and *ncan* are primarily expressed in the nervous system [20, 26, 39–49], though notochord and heart expression has also been reported [46, 50–52]. Of the four *lectican*s, mutation of *acan* leads to the most significant defects, including severe chondrodysplasia [23], while *vcan* loss-of-function causes abnormal eye and heart development [53–55]. The functions of *bcan* and *ncan* are less clear, however, as mice deficient in these genes show only minor defects in neuronal potentiation [56, 57].

It has been proposed that the evolution of novel interactions between Lecticans, hyaluronan, and other glycoproteins played an important role in the evolution of vertebrate tissues [19]. However, our understanding of *lectican* expression, function, and evolution is based entirely on the information from model gnathostomes. It is thus unclear when in the vertebrate lineage *lecticans* originated, were duplicated, and acquired their diverse functions. The only two living jawless vertebrates, the lampreys and hagfish (the cyclostomes) have been indispensable for understanding vertebrate evolution [58–62]. These modern agnathans diverged from the lineage leading to gnathostomes around 500 million years ago [63, 64]. Due to accessibility, lampreys are the best studied of the two, and historical and modern molecular comparisons have shown that lamprey and gnathostomes share many core aspects of their development[58, 65–69].

In this study, we used genomic and transcriptomic data from the sea lamprey, *Petromyzon marinus* to gain insight into the evolutionary history of *lectican* genes. These data support independent expansion of the *lectican* family in the lamprey and gnathostome lineages. We also characterized the expression patterns of *lecticans* in sea lamprey embryos and larvae, and show that *lectican* expression in neural, cardiac, and skeletal tissue is highly conserved across living vertebrates. In contrast, we find that expression of *lectican* paralogs in the head skeleton is markedly different between lamprey and gnathostomes. We posit that the ancestral pre-duplication *lectican* had a complex expression pattern which was independently partitioned between paralogs in the lamprey and gnathostome lineages. We further speculate that the primordial Lectican protein functioned to facilitate mesenchymal histology and behavior in the first vertebrates.

## RESULTS

### The sea lamprey has four *lectican* genes encoding proteins with similar domain structures

We searched the *P. marinus* germline genome [70] and identified four different genomic scaffolds containing exons with sequence similarity to gnathostome Lecticans. We also searched all publicly available lamprey transcriptome data, as well as our own database of transcriptome sequences (see Methods) for gnathostome *lectican*-like sequences, and assembled these into 4 mRNAs corresponding to proteins of 1871aa, 1757aa, 1825aa, and 1343aa respectively (see Tab. S3 for accession numbers). All identified *lectican* exons aligned to parts of the reconstructed mRNAs, indicating there are only four sea lamprey *lectican* genes. We named these genes *lecticanA* (*lecA*), *lecticanB* (*lecB*), *lecticanC* (*lecC*), and *lecticanD* (*lecD*). We then searched for conserved domains in lamprey *lectican* conceptual translation products using NCBI’s Conserved Domain search tool, and by alignment with gnathostome Lecticans. We found that although all lamprey Lectican protein sequences had largely archetypical domain structures, at least one domain appeared to be missing in each [Fig. 1A]. LECA and LECC did not possess an identifiable complement control protein domain, while LECB did not have an immunoglobulin-like domain, and LECD did not have EGF-like domains. We also found that no lamprey Lectican possessed the extra X-link domain seen in ACAN [Fig. 1A].

**Figure 1.**
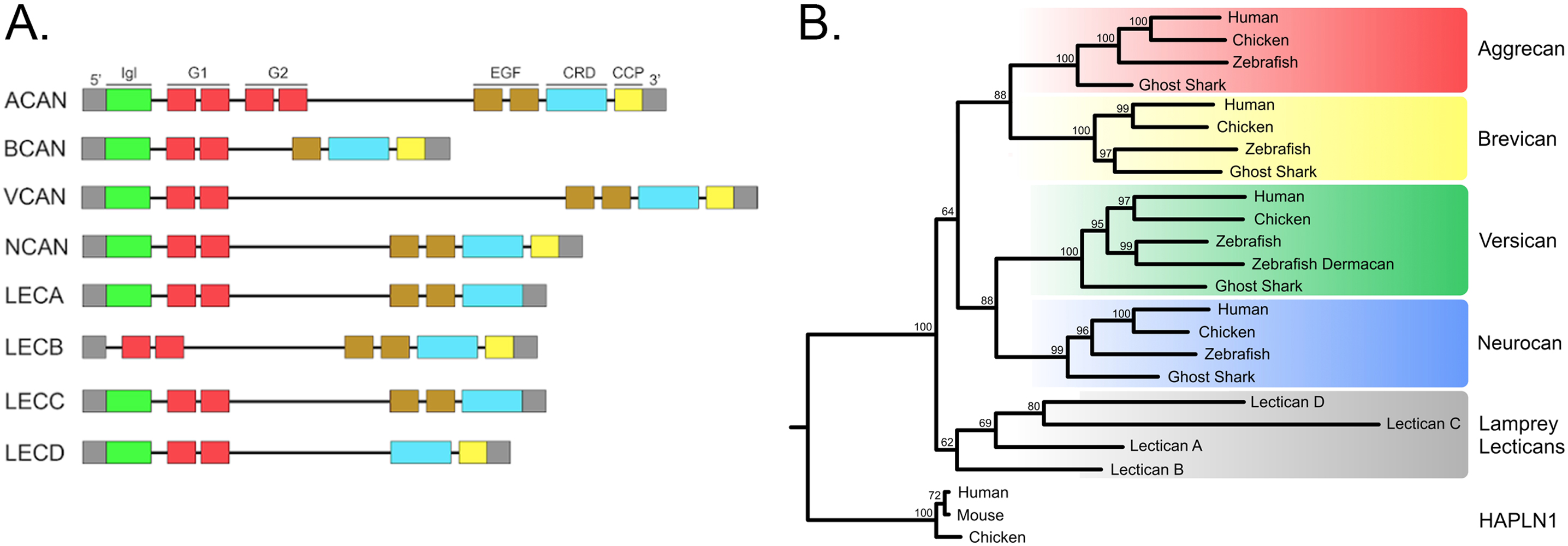
Structural comparison and molecular phylogeny of vertebrate lecticans. (A) Domain structure of vertebrate Lecticans with the N-terminus to the left. **Keywords:** Igl: Immunoglobulin- like domain; G1 / G2: link domains; EGF: EGF-like domains; CRD: carbohydrate recognition domains; CCP: complement control protein domain. (B) Phylogenetic relationships of vertebrate Lecticans based on amino acid sequence alignments. Lamprey sequences are in gray boxes while individual gnathostome paralogy groups are in colored boxes. Maximum likelihood analysis scores are shown at the respective node. HAPLN1 sequences were designated as outgroup. Original tree and accession numbers for all sequences can be found in Fig. S1 and Tab. S1.

### Phylogenetic analyses do not support one-to-one orthology of lamprey and gnathostome *lectican*s

To deduce relationships between lamprey and gnathostome *lectican*s, we used *lectican* protein sequences to perform maximum likelihood phylogenetic analyses, with different taxa, substitution models, and individual parameters for tests [71–74] [Fig. S1,S2,S3,S4]. Among gnathostome *lectican*s, we recovered all four known paralog groups and found good support for *acan*+*bcan* and *vcan*+*ncan* subfamilies. In contrast, we found that none of the lamprey Lecticans consistently group within any of the four gnathostome Lectican paralogy groups, nor the *acan*+*bcan* and *vcan*+*ncan* subfamilies regardless of the parameters used to build the phylogenies [Fig. 1B, Fig. S5]. Lamprey *lecticans* and *hapln*s likely originated from a tandem duplication event early in the vertebrate lineage. We reasoned that building a phylogenetic tree using HAPLNs and the HAPLN-aligning portion of Lectican protein sequences might help resolve the relationships between lamprey and gnathostome Lecticans [Fig S4]. As with the full-length Lectican phylogeny, none of the lamprey Lecticans grouped with any gnathostome paralogy group with high confidence.

### Analyses of syntenic genes also fails to conclusively support one-to-one orthology between lamprey and gnathostome *lectican* paralogs

All gnathostome *lectican* paralogs are adjacent to a corresponding HAPLN paralog [18]. We thus searched for HAPLN-like reading frames in the lamprey genome [70], and used these to create a phylogeny of chordate HAPLN-related genes in hopes of resolving the relationships between vertebrate *lectican/HAPLN* loci. We identified one lamprey *HAPLN* gene linked to *lecticanD*. However, as with lamprey Lecticans, lamprey HAPLN fails to group convincingly with any single gnathostome paralogy group [Fig. S6]. We also found that gnathostome HAPLN1s and HAPLN4s form a weakly supported clade, consistent with the relationships of their adjacent *lecticans, vcan* and *ncan*. We expanded our search to include other possible conserved syntelogs. We found that all gnathostome and lamprey *lecticans* are linked to paralogs of the myocyte enhancer factor *mef2* gene family. We thus created a phylogenetic tree of MEF2 amino acid sequences to see if it could provide insights into the evolution of the vertebrate *lectican* locus. As with HAPLN genes, none of the lamprey MEF2 sequences clustered convincingly with any gnathostome MEF2 paralogy group using any parameters [Fig. S7].

As a final test of orthology between lamprey and gnathsotome *lectican*s, we compared the gene complement around the gnathostome and lamprey *lectican* loci. For each lamprey *lectican*, we asked if any of the surrounding 40 genes (when available) had homologs that were syntenic with any chick, spotted gar, or elephant shark *lecticans* [Fig. 2A, Fig. S8]. We found that *lecA* had the most conserved syntelogs, with 21/40 of adjacent genes having gnathostome homologs closely linked to one or more *lecticans* (i.e. syntelogs). Of those, 15 were exclusively linked to an *acan* or a *bcan*, while only 4 were exclusively linked to a *vcan* or *ncan*. Around the *lecB*, *lecC*, and *lecD* loci, 30-40% of genes had unambiguous gnathostome syntelogs, with similar proportions linked to the *acan*+*bcan* versus *vcan*+*ncan* subfamilies [Fig. S8]. Thus, comparisons of syntelogs provide some support for placing *lecA* in the *acan*+*bcan* subfamily [Fig 2B].

**Figure 2.**
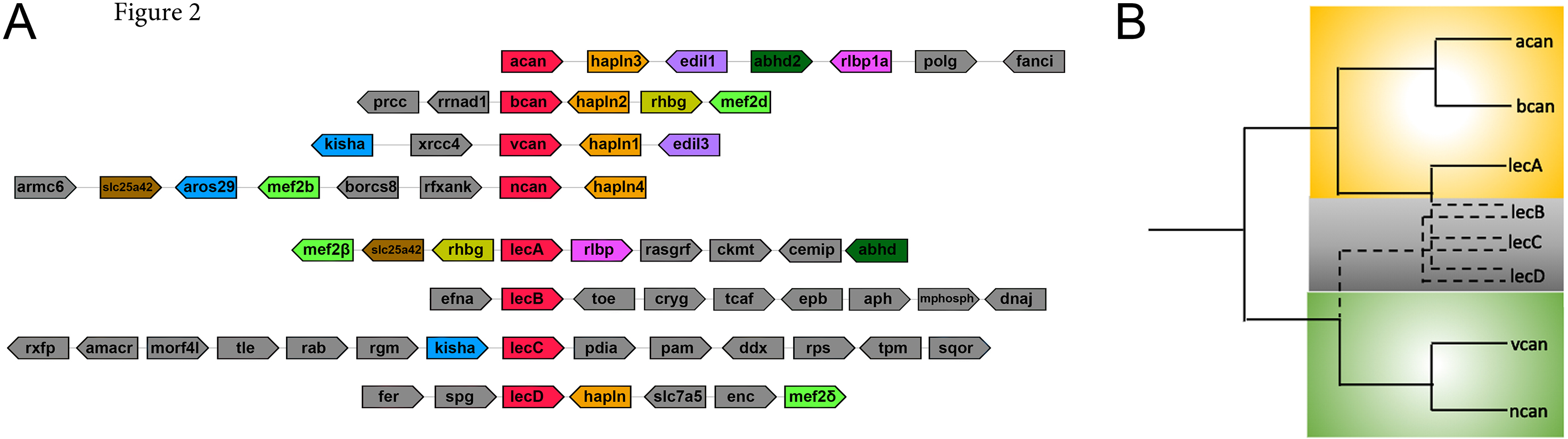
Summary of *lectican* microsynteny and implications for their evolutionary relationships. (A) Conserved genes adjacent to the gnathostome (top four) and lamprey (bottom four) lectican loci. Syntelogs are shown in orientation with respect to their linked *lectican* gene. *Lectican*s are in red. Homologous genes are colored the same. Gnathostome genes were surveyed within a 300kb radius while lamprey genes were surveyed within a 400kb radius. Macrosyntenic analyses can be found in Table S1. (B) Hypothetical scenarios for the evolution of vertebrate lecticans when synteny data is incorporated. While syntenic analyses suggest that *lecticanA* is orthologous to the *aggrecan*/*brevican* gene family, the relationship of *lecticanB*, *lecticanC*, or *lecticanD* is unclear.

### Expression of *lecticans* in sea lamprey embryos and larvae

We first detected *lecA* expression at Tahara [75] stage 21 (st. T21) in the presumptive neural tube and newly formed somites [Fig. 3A]. At st. T23, we continued to see lecA expression in these regions, and sectioning revealed transcripts in the notochord, neural tube floor plate, and sclerotome [Fig. 3C, 3C’]. *lecA* transcripts were also detected in the developing myocardium at this stage. By T24 and T25, *lecA* expression expanded into the posterior lateral line ganglia, zona limitans intrathalamica, and the telencephalon [Fig. 3D, D’, D’’]. At stage T26, we observed new expression in the posterior heart tube [Fig 3E]. At this stage, *lecA* transcripts were also found in skeletogenic mesenchyme in the pharynx and oral region, and in the fin fold mesenchyme [Fig 3E’, E’’]. We also noted that expression of *lecA* in the maturing pharyngeal arches was highly dynamic, with activation and downregulation occurring in an anterior to posterior wave. By st. T27, this pharyngeal mesenchyme cell expression was limited to the oral hood, outer velum, and lips [Fig 3F]. At stage T28, *lecA* was almost entirely restricted to the mucocartilage of the oral hood, velum, and fin fold [Fig 3G, G’, 3I, I’].

**Figure 3.**
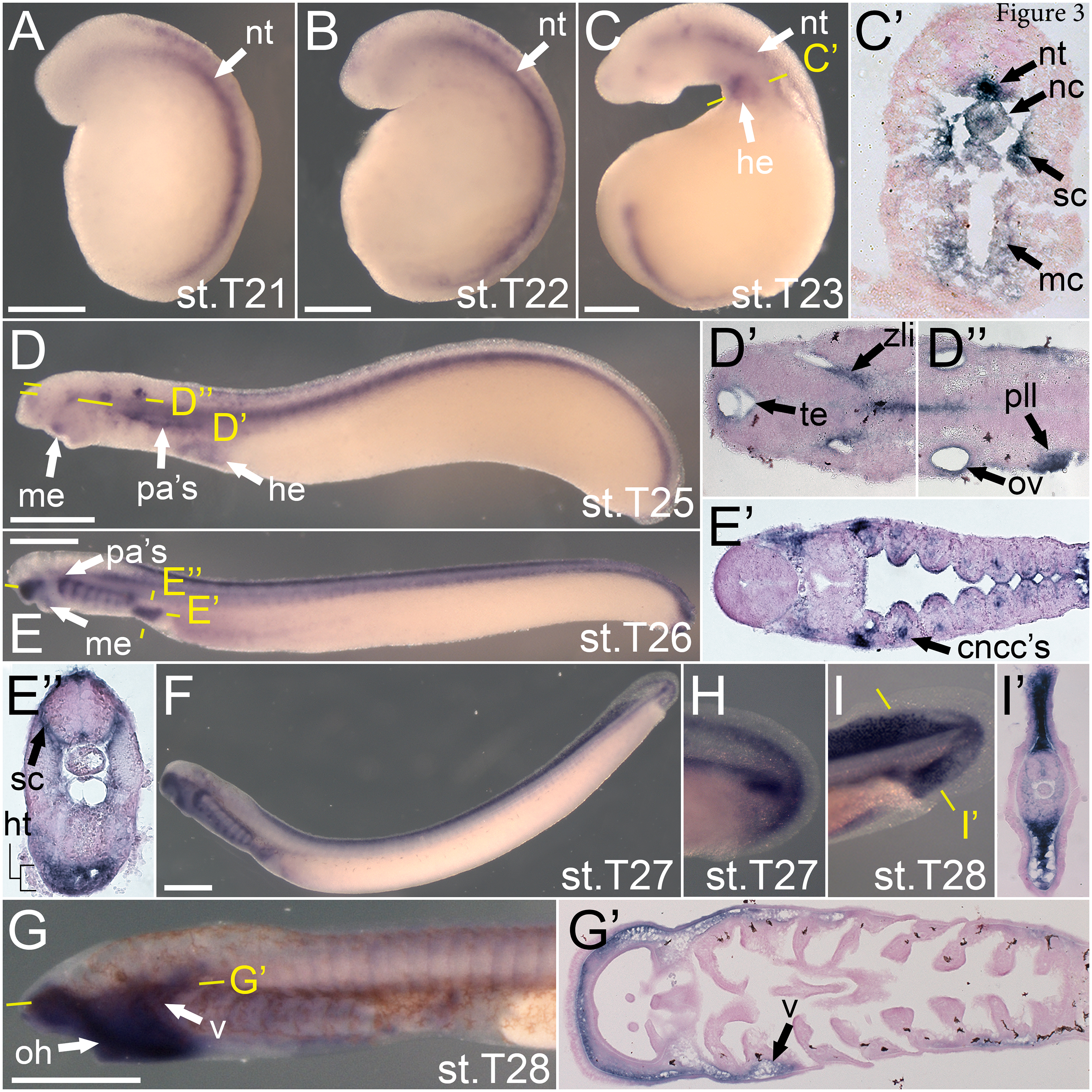
Expression of *lecA* in embryos and larvae. Left lateral view in all non-prime panels. Developmental stage (Tahara, 1988) for each whole mount panel is in the bottom right corner. Prime panel stages correspond to their whole mount. For all non-prime panels, scale bar represents 500 μm (A-B) *lecA* is expressed in the neural tube and presumptive somites (C-C’) *lecA* is expressed in the myocardium of the heart, the notochord, neural tube floor plate, and medial sclerotome. (D,D’,D”) *lecA* expression is in the pre-oral mesenchyme, the dorsal aspect of the nascent pharyngeal arches, the otic vesicle, and the developing heart and brain (E, E’, E”) *lecA* is expressed in the the premandibular mesenchyme, the dorsal sclerotome lateral to the neural tube, pharyngeal neural crest cells, and developing heart tube. (F, H) *lecA* is localized in the developing head skeleton as well as fin mesenchyme, but is absent from the brain by this stage. (G, I) *lecA* expression is in the oral hood and velar mucocartilage as well as fin mesenchyme. (H-I) Focused view of *lecA* expression in the developing fin. **Keywords:** cncc’s: cranial neural crest cells; he: heart; ht: heart tube; mc: myocardium; me: pre-oral mesenchyme; nc: notochord; nt: neural tube; oh: oral hood; ov: otic vesicle; pa’s: pharyngeal arches; pll: posterior lateral line ganglion; sc: sclerotome; te: telencephalon; v: velum; zli: zona limitans intrathalamica

Expression of *lecB* was first observed at stage T23 in the oral ectoderm [Fig 4A]. This expression remained similar until mid pharyngula at (T25) when *lecB* expression expanded into the lateral neural tube [Fig 4B, B’]. By middle-late pharyngula at stage T25 and T26, *lecB* was observed in the pronephros [Fig 4C, 4D]. We also identified *lecB* expression in the nasohypophyseal and ophthalmic, lens, and maxillomandibular placodes as well as the basolateral hypothalamus [Fig 4D’]. As skeletogenesis began at stages T26.5 and T27, we identified *lecB* transcripts in the mucocartilage of the outer velum, lips, and ventrolateral pharyngeal bars [Fig 4E, E’]. At these stages, we were able to confirm *lecB* expression in the pharyngeal endoderm through sectioning. However, by stage T27, this expression began to fade in an anterior-posterior manner. We no longer detected *lecB* in the developing brain or neural placodes likewise at these stages. By stage T28, *lecB* was primarily found in the medioventral cartilage bar and the developing oral papillae [Fig 4F, F’].

**Figure 4.**
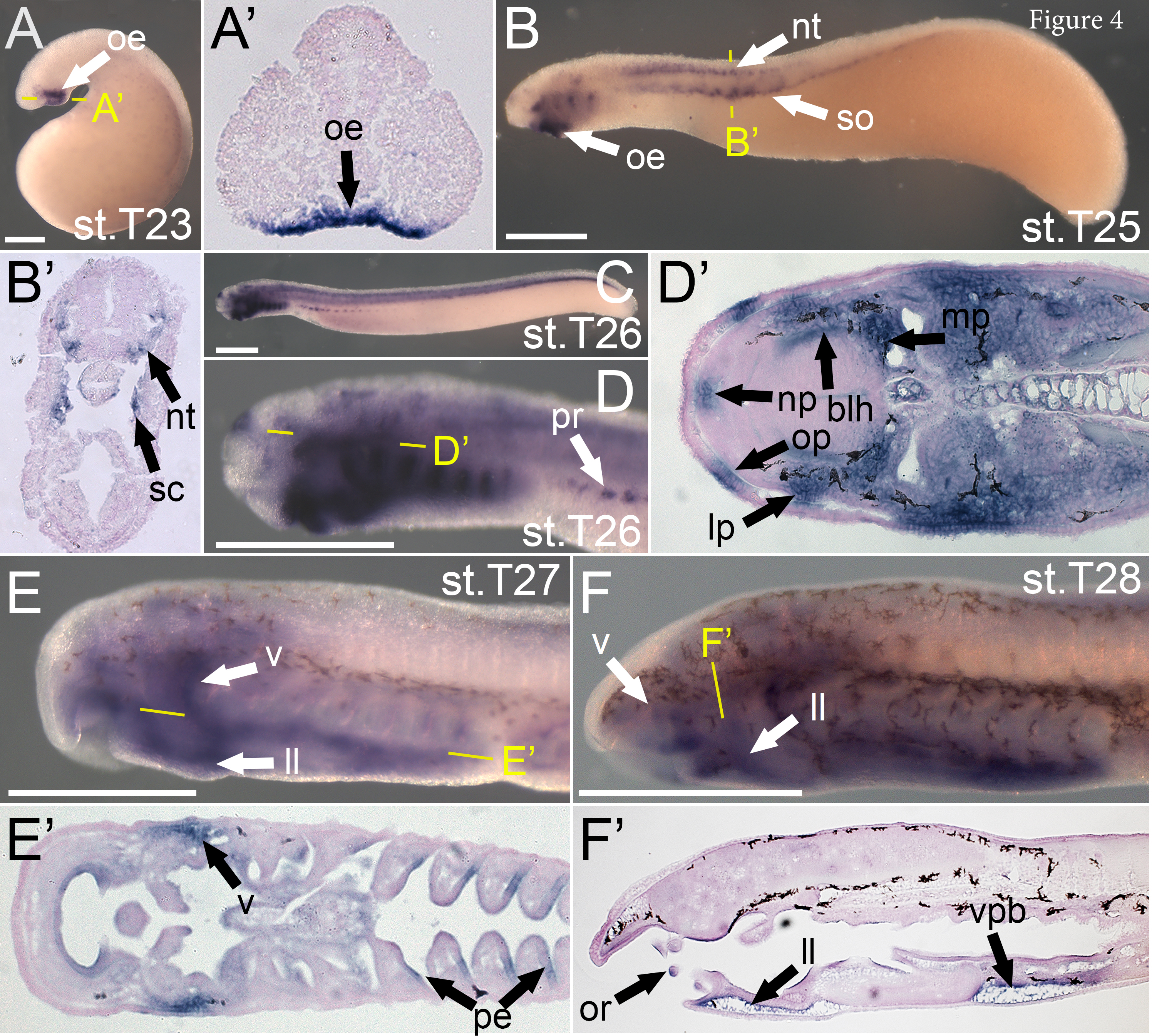
Expression summary for *lecticanB* in *P. marinus* embryos. Left lateral view in all non- prime panels. Developmental stage (Tahara, 1988) for each whole mount panel is in the bottom right corner. Prime panel stages correspond to their whole mount. For all non-prime panels, scale bar represents 500 μm. (A-A’) *lecB* expression is in the oral ectoderm (B-B’) *lecB* expression is in the oral ectoderm, lateral neural tube, and medial sclerotome (C,D,D’) *lecB* expression is in the developing brain, neural placodes, and pronephros. (E,E’) *lecB* expression is in the pharyngeal endoderm as well as the mucocartilage of the lateral velum and lower lip. (F,F’) *lecB* expression is in the mucocartilage of the ventrolateral pharyngeal bars and lower lip as well as the oral papillae. **Keywords:** blh: basolateral hypothalamus; ll: lower lip; lp: lens placode; mp: maxillomandibular placode; np: nasohypophyseal placode; nt: neural tube; oe: oral ectoderm; op: ophthalmic placode; or: oral papillae; pe: pharyngeal endoderm; pr: pronephros; sc: sclerotome; so: somites; v: velum; vpb: ventrolateral pharyngeal bars

Strikingly, *lecC* expression was only observed in forming cell-rich hyaline cartilage bars in the head skeleton. This expression closely tracks alcian blue reactivity, as previously described [76]. We first detected *lecC* transcripts at stage T26.5 in neural crest in the intermediate domain of the third through sixth pharyngeal arches [Fig 5A]. This expression expanded to the seventh and eighth arch cartilage bars, and by stage T28, *lecC* expression was seen in all hyaline cartilage bars in the posterior pharynx, as well as the trabeculae [Fig 5C, C’].

**Figure 5.**
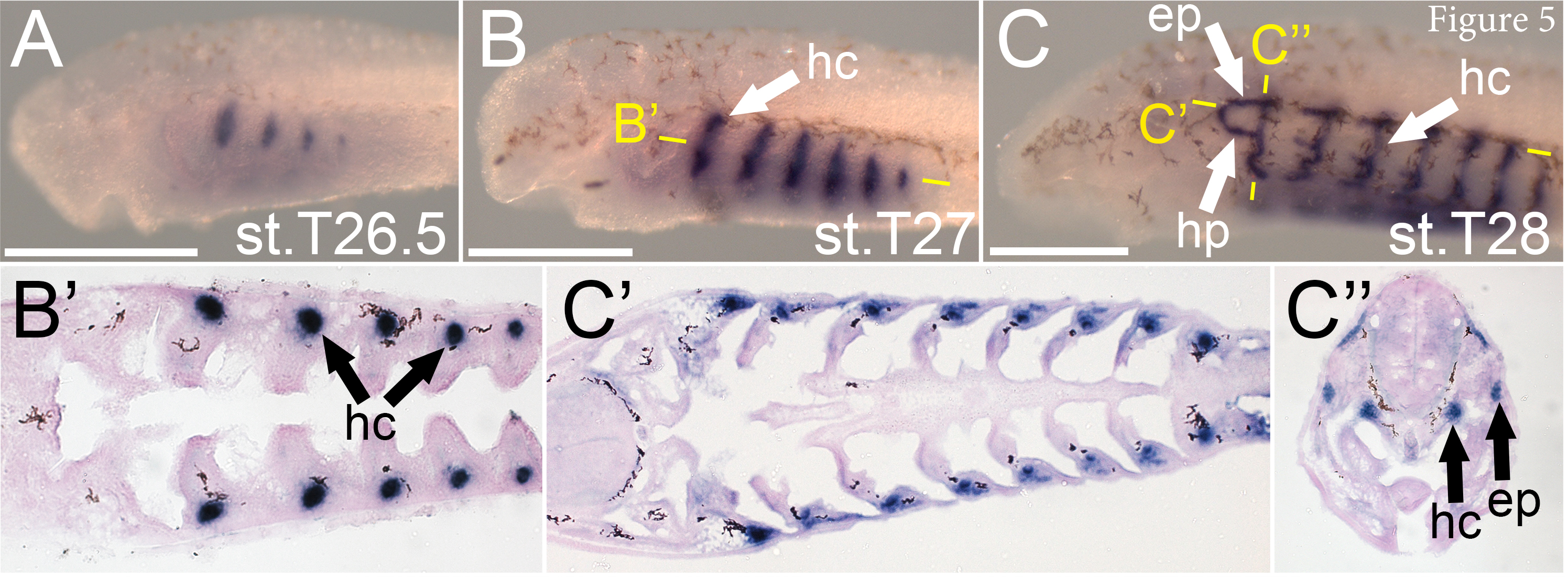
Expression summary for *lecticanC* in *P. marinus* embryos. Left lateral view in all non- prime panels. Developmental stage (Tahara, 1988) for each whole mount panel is in the bottom right corner. Prime panel stages correspond to their whole mount. For all non-prime panels, scale bar represents 500 μm. (A-B) *lecC* is found in the cell-rich hyaline cartilage of the pharynx. (C) *lecC* is in the pharyngeal hyaline cartilage, the epitrematic and hypotrematic processes of the pharynx, as well as the trabeculae. **Keywords:** ep: epitrematic process; hc: hyaline cartilage; hp: hypotrematic process

We identified *lecD* expression at stage T21 in the developing somites [Fig 6A]. By early pharyngula in stage T23, *lecD* was additionally found in the splanchnic mesoderm [Fig 6B, B’]. At this stage, our sectioning confirmed expression in the somites to be localized in the sclerotome [Fig 6B’]. Expression in the somite abated by stages T24 and T25, starting in the anterior somites and moving posterior [Fig 4C]. At stage 26, we detected *lecD* in the posterior endocardium and heart tube as well as the ventral aorta [Fig 4D, 4D’]. By late pharyngula at stages T26.5 through T27, *lecD* was expressed in the entirety of the aortic arches in the pharynx as well as the mucocartilage of the lower lip and ventral pharynx surrounding the endostyle [Fig 4E, F, F’, F’’]. Expression in the aortic arches and ventral mucocartilage dissipated by stage T28, but we continued to see expression in the heart tube, lower lip mucocartilage, and ventral aorta [Fig 4G, G’ G’’].

**Figure 6.**
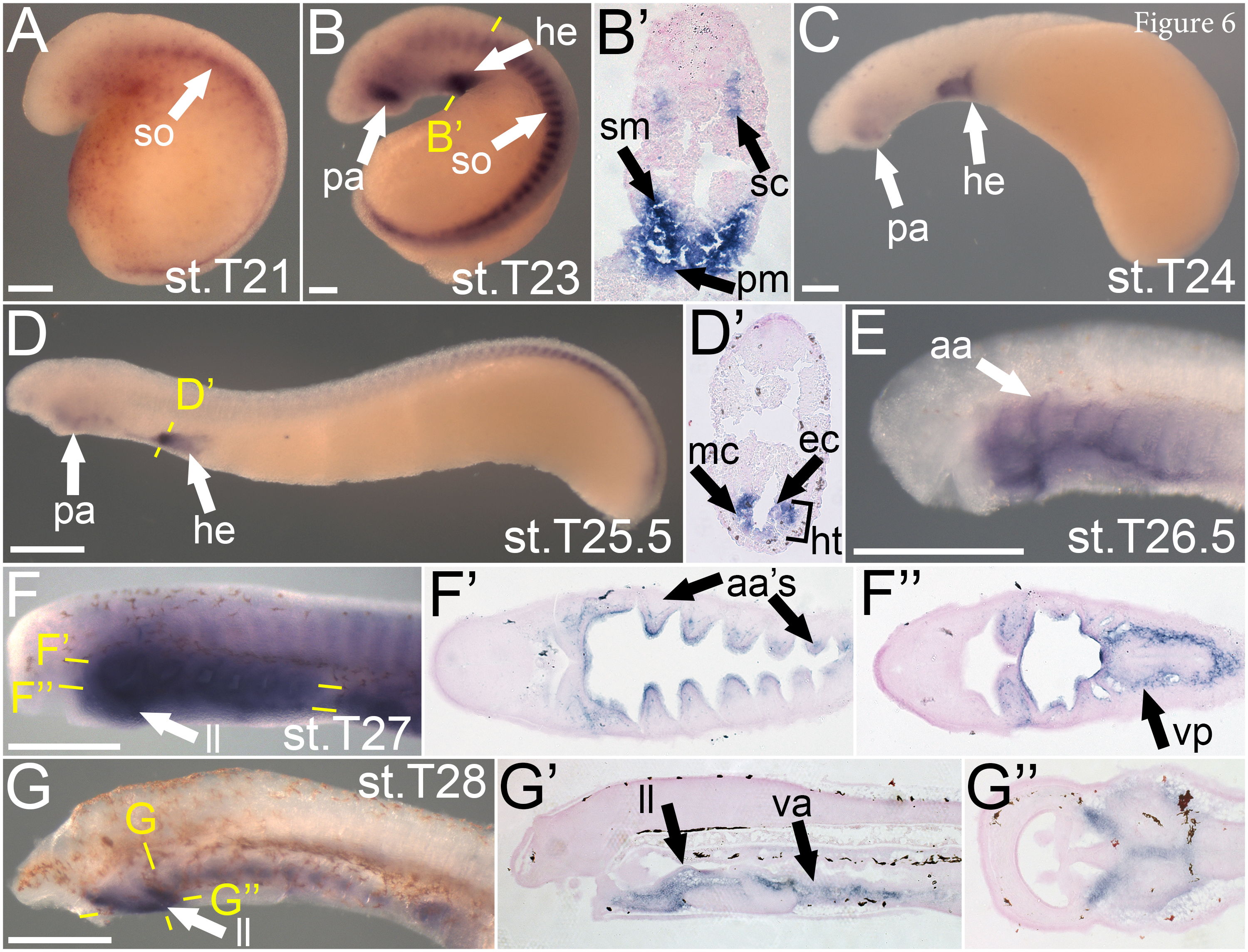
Expression summary for *lecticanD* in *P. marinus* embryos. Left lateral view in all non-prime panels. Developmental stage (Tahara, 1988) for each whole mount panel is in the bottom right corner. Prime panel stages correspond to their whole mount. For panels A-C, scale bar represents 250 μm. For panels D-G, scale bar represents 500 μm (A) *lecD* is found in the somites. (B-B’) *lecD* is additionally found in the first pharyngeal arch as well as the sclerotome and developing splanchnic mesoderm of the heart. (C) *lecD* expression ablates in the somites but continues in the first pharyngeal arch as well as the developing heart. (D-D’) *lecD* is localized in the first pharyngeal arch, the heart tube, endocardium, and splanchnic mesoderm. (E) *lecD* is expressed in the developing aortic arches. (F) *lecD* expression is in the pharyngeal vasculature as well as the mucocartilage of the ventral pharynx. (G) lecD is expressed in the ventral aorta as well as the mucocartilage of the lower lip. **Keywords:** aa: aortic arches; ec: endocardium; he: heart; ht: heart tube; ll: lower lip mucocartilage; pa: first pharyngeal arch; sc: sclerotome; sm: splanchnic mesoderm; so: somites; va: ventral aorta; vp: ventral pharynx mucocartilage

## DISCUSSION

The evolution of vertebrate developmental and morphological novelties has been linked to a variety of genetic and genomic events, including the evolution of new gene regulatory interactions between ancient developmental regulators, the origin of new gene families, and genome-wide duplication events [5, 7, 13, 15, 77]. To better understand the role of new gene families and gene duplications in vertebrate morphological evolution, we investigated the phylogeny and expression of *lectican* genes in the sea lamprey. To our knowledge, this work constitutes the first comprehensive expression analysis of all *lecticans* in a single vertebrate, and the first description of these genes in a jawless vertebrate.

### Evolutionary history of the *lectican* family

Gnathostome *lectican*s and the related *hapln*s are thought to have arisen via tandem duplication of a *hapln*-like gene sometime in the vertebrate lineage, with *lectican*s later gaining their complex domain structure by exon shuffling. Consistent with this scenario, we found one lamprey *hapln* closely linked to the *lecD* locus [Fig. 2A]. The timing of the duplication events that created the gnathostome and lamprey *lectican* families is less clear. Like previous reports, our phylogenetic analyses place all gnathostome *lecticans* into four paralogy groups, with *acan*+*bcan* and *vcan*+*ncan* forming two subfamilies. This topology is typical of gnathostome gene families and strongly suggests the four gnathostome *lecticans* were generated during the two vertebrate genome-scale duplication events (1R and 2R) [4, 78–80]. In contrast, the relationships among lamprey *lecticans*, and between lamprey and gnathostome *lecticans*, are inconclusive. Regardless of tree building parameters, lamprey *lectican*s fail to consistently group with gnathostome paralogy groups, often clustering weakly with each other [Fig. 1B].

Phylogenetic analyses of the neighboring genes *hapln* and *mef2,* and comparisons of syntenic genes yielded similarly inconclusive and weakly-supported phylogenies. There are several scenarios that could account for lack of clear one-to-one orthology between lamprey and gnathostome *lectican*s. One explanation is that the lamprey and gnathostome *lectican*s are the result of independent duplications of a single ancestral *lectican* in each lineage. At the other extreme, lamprey *lectican*s could be fast-evolving cryptic orthologs of gnathostome *lectican*s generated by the two vertebrate genome-wide (2R) duplication events. Various scenarios involving shared duplication, gene loss, and independent duplication are also possible. A prerequisite for cryptic one-to-one orthology is that lamprey diverged from gnathostomes after the 2R genome duplications. However, recent comprehensive comparisons of chordate genome structure refute this, showing lamprey most likely diverged from gnathostomes before the second, “2R”, genome duplication [4, 81]. If this is the case, the common ancestor of lamprey and gnathostomes likely had two *lectican*s, an ancestral *acan+bcan* and an ancestral *vcan*+*ncan*. Consistent with this, we find that the genomic region surrounding *lecA* is *acan*+*bcan*-like but shows no particular similarity to either the *acan* or *bcan* regions (Tab. S1). In contrast, we find that none of the other lamprey *lectican* genomic regions are *ncan* and/or *vcan*-like. Taken together, our data support the presence of two *lectican*s in the last common ancestor of lamprey and gnathostomes, with one or both being duplicated in the cyclostome lineage to yield the four lamprey *lectican*s [Fig. 2B]. Of these, *lecA* is likely derived from the *acan*+*bcan*-related 1R duplicate, while the orthology of *lecB*, *lecC*, and *lecD* and gnathostome *lectican*s is unresolved [Fig. 2B]. Although *lecB*, *lecC* and/or *lecD* could be cryptic *vcan*+*ncan* family members, it is also possible the *vcan*+*ncan* subfamily was lost in the lamprey lineage, and all lamprey *lectican*s are *acan*+*bcan* co-orthologs [Fig. 2B].

### Lectican expression in the nervous system is ancestral within vertebrates

Regardless of their phylogenetic relationships, we found that almost every gnathostome *lectican* expression domain was conserved in lamprey, with only a few minor differences. To what degree these differences reflect divergence in *lectican* regulation between lamprey and gnathostomes, or the incomplete documentation of *lectican* expression in gnathostomes, is unclear. In the central nervous system, *lecA* and frog *vcan* both display expression in the neural tube floor plate [Fig 3C’] [30]. Lamprey *lecB* expression is also observed in the lateral neural tube [Fig 4B’], though there are no reports of gnathostome *lectican* transcription in this region. *lecA* and *lecB* are expressed in the developing brain like *bcan* and *ncan* [Fig 3D’, D’’, Fig 4D’], though not as broadly. Like *ncan*, *bcan*, and *vcan, lecA* and *lecB* are expressed in the cranial placodes and sensory ganglia [Fig 3D’, D’’, Fig 4D’], though in different neural populations [38, 46, 47]. Although *lectican* expression in the forming nervous system appears to conserved among living vertebrates, the role of *lecticans* in neural development is unclear, as *bcan* and *ncan*-deficient mice show only minor differences in neuron function [56, 57]. Regardless of their precise functions, our data suggest that the LCA of cyclostomes and gnathostomes expressed *lecticans* in both the peripheral and central nervous systems.

### Lectican expression in mesoderm-derived tissues is conserved across vertebrates

As in the forming nervous system *lectican* expression in mesodermal derivatives is largely conserved between lamprey and gnathostomes. In the gnathostome heart, *aggrecan* marks migratory cardiac mesoderm [29], while *ncan* marks the forming myocardium and splanchnic mesoderm, and *vcan* marks the endocardium and the heart tube [29, 46, 51, 82]. In lamprey, *lecA* is expressed in the myocardium [Fig 3E’’] while *lecD* marks the posterior endocardium and heart tube as well as the ventral aorta [Fig 6D’]. As in neural tissue, the precise role of *lectican*s in the gnathostome heart is unclear, though mouse *vcan* mutants have major defects in the developing heart tube and endocardial cushion [53–55].

Aside from cardiac mesoderm, we also noted expression of one or more lamprey *lectican*s in the notochord [Fig 3C’], pronephros [Fig 4C], fin mesenchyme [Fig 3I], and sclerotome [Fig 3C1’, 4B’, 6B’]. All of these mesodermal tissues express one or more *lectican*s in gnathostomes in temporal and spatial patterns virtually identical to their lamprey counterparts.

The only notable difference in mesodermal *lectican* expression we observed was an absence of lamprey *lectican*s in somatic lateral plate mesoderm (LPM), which gives rise to *acan* and vcan- expressing skeletal tissue in gnathostome paired fins and limbs.

### Combinatorial *lectican* expression suggests lamprey possesses a diverse array of neural crest-derived skeletal tissues

We find that expression of multiple *lectican*s in forming and differentiated skeletal tissue is a conserved feature of vertebrate development. However, we also noted that gnathostomes typically transcribe only two *lectican*s in skeletogenic neural crest cells, *acan* and *vcan*, whereas lamprey expresses all four. Furthermore, lamprey *lecticans* are expressed in spatiotemporally distinct patterns throughout development, creating a combinatorial code of *lectican* expression in different parts of the nascent lamprey head skeleton. The histological heterogeneity of the lamprey head skeleton, which includes a mesenchymal chondroid tissue called mucocartilage, has been noted before [83–89]. Anatomical work on adult hagfishes has also revealed diverse histology in the head skeleton [90–93], suggesting that the LCA of cyclostomes likely had multiple chondroid tissue types. It is possible the combinatorial co-expression of *lectican*s in the lamprey head skeleton elements reflects histological differences between different subtypes of mucocartilage. If this is the case, it would suggest that either 1) the LCA of cyclostomes and gnathostomes had a diversity of neural crest-derived chondroid tissues and the gnathostome lineage has retained only a few; or 2) the LCA of cyclostomes and gnathostomes had only a few neural crest-derived cartilage subtypes and the diversity seen in the sea lamprey head skeleton is a derived feature of lampreys, or cyclostomes. It has been previously shown that the pharyngeal skeleton of cyclostomes is patterned using the same basic mechanisms as seen in gnathostomes [9, 66, 88, 94–97] . In gnathostomes, this patterning acts a scaffold for proper deployment of the morphogenetic programs that control skeletal element shape and the tissue differentiation. In lamprey, which has a largely symmetrical oropharygneal skeleton, this patterning may function mainly to control the activation of distinct differentiation programs in different parts of the head skeleton as previously proposed [66, 95, 96].

### Different patterns of specialization and subfunctionalization after *lectican* duplication in lamprey and gnathostomes

Gene duplication is thought to facilitate evolutionary novelty by creating additional copies of genes that can then diverge to gain new expression domains and functions (neofunctionalization). More commonly, however, duplication leads to partitioning of ancestral expression domains (subfunctionalization) as described by the duplication-degeneration-complementation model [98]. Recent functional genomic comparisons have also highlighted the importance of specialization after duplication in the vertebrate lineage. During specialization, one paralog loses most aspects of its ancestral expression pattern and becomes specialized for a particular domain, while other paralogs maintain the complete ancestral pattern [12]. Our data suggest the ancestral *lectican* had a complex expression pattern, and was independently duplicated in the lamprey and gnathostomes, with little apparent neofunctionalization in either lineage [Fig. 7]. We also find that the relative roles of specialization and subfunctionalization differ between the gnathostome and lamprey *lectican* families. Striking specialization is apparent in the lamprey *lectican* family, where *lecA* is transcribed in virtually all major *lectican* expression domains, while *lecC* has highly restricted expression in cell-rich hyaline cartilage [Fig. 7]. Similarly, *lecD* transcripts are only seen in the sclerotome, heart, and a subpopulation of skeletogenic NCCs. In contrast, no gnathostome *lectican* is so strictly specialized, and all paralogs are expressed in partially overlapping subsets of the ancestral expression pattern that could be described as “overlapping subfunctionalization” [22, 23, 25, 29, 33, 34, 39, 44, 45, 50] [Fig. 7]. Whether the different modes of expression pattern evolution have any significant consequences for Lectican protein function is unclear. Specialization of ohnologs is usually associated with rapid divergence in protein coding sequence [12]. Contrary to this prediction, lamprey *lecC*, the most specialized lamprey *lectican*, and *lecA*, the most broadly expressed lamprey *lectican* have similar, archetypical *lectican* structures. Meanwhile, all gnathostome *lectican*s vary significantly in length, and have lost and gained different functional domains.

**Figure 7.**
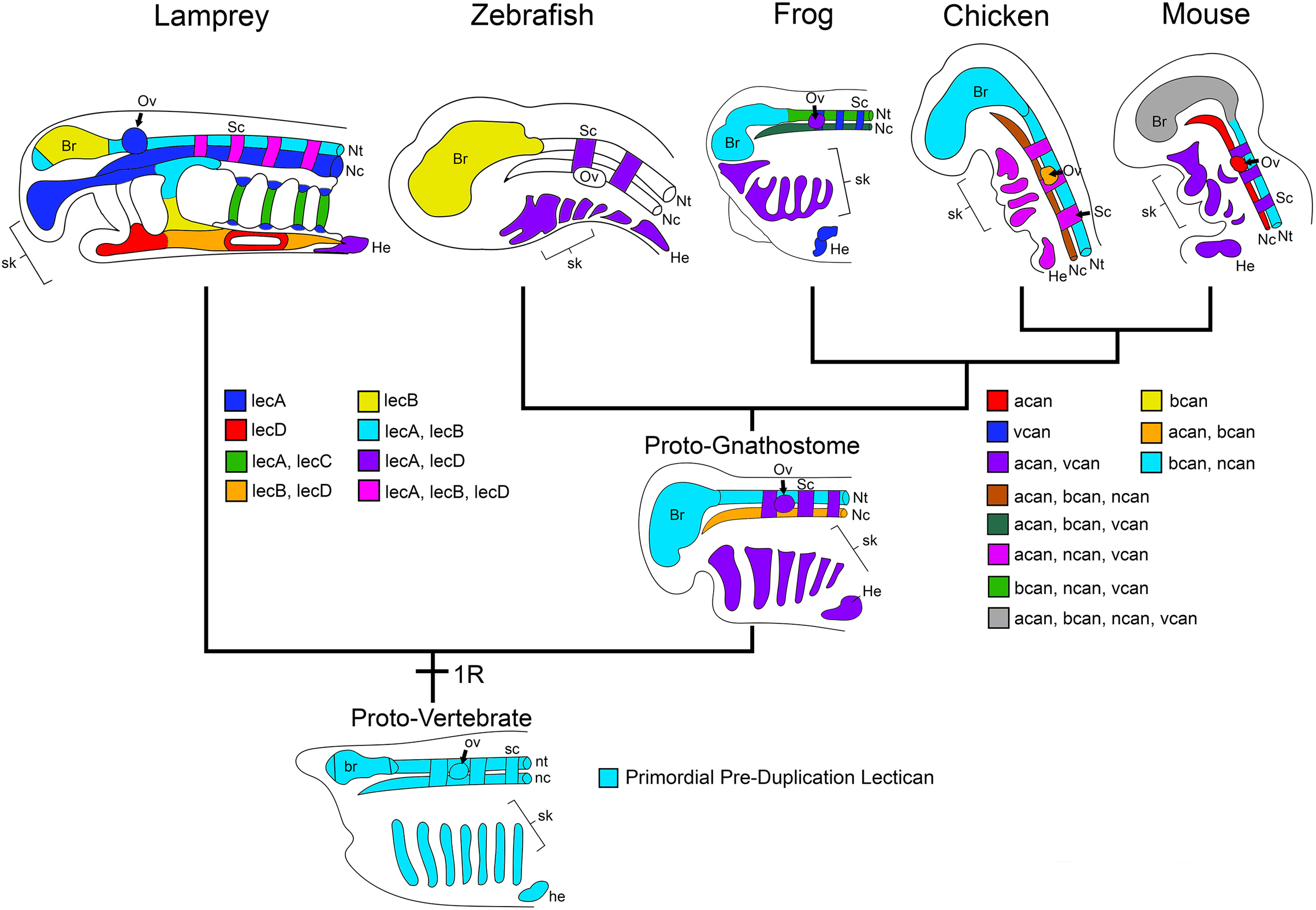
The evolution of *lectican* expression patterns in the head. Modern gnathostome lectican expression is depicted based on current data for zebrafish (*Danio rerio*), frog (*Xenopus laevis*), chicken (*Gallus gallus*), and mouse (*Mus musculus*), and their “average” is depicted as an idealized proto-gnathostome. Expression data for chondrichthyans is currently not available. **Keywords:** br: brain; he: heart; nc: notochord; nt: neural tube; ov: otic vesicle; sc: sclerotome; sk: skeletal mesenchyme

Nevertheless, it is provocative that both lamprey and gnathostomes typically express multiple *lecticans* in each expression domain. This suggests Lectican proteins are not entirely redundant, and supports the idea that combinations of functionally distinct Lectican proteins may confer subtle histological differences in related tissues.

### The primordial *lectican* likely contributed to the evolution of vertebrate traits and functioned to support mesenchymal histology

Vertebrate evolution involved the acquisition of new organs, tissues, and cell types, as well as the elaboration of many pre-existing cell and tissue types. The *lectican* family is also a vertebrate novelty that arose at around the same time as these histological and morphological innovations. To what degree the evolution of new gene families drove the evolution of vertebrate traits is an open question. We used our expression data to ask if *lectican*- expressing cells and tissues were usually vertebrate novelties, or had clear homologs in invertebrate chordates [Tab. 1]. If a recognized homolog was present, we next asked if the histology of the vertebrate cell/tissue differed fundamentally from its invertebrate counterpart.

**Table 1.** Lecticans are mainly expressed in cells and tissues that are unique to vertebrates, or have distinct histology in vertebrates.

|  | Anterior Mesoderm and/or Cranial Neural Crest |  |  | Axial Mesoderm | Paraxial Mesoderm |  | Intermed. Meso. | Lateral plate Mesoderm |  | Oropharyngeal Epithelia |  | Neural |  |
| --- | --- | --- | --- | --- | --- | --- | --- | --- | --- | --- | --- | --- | --- |
|  | Cell-rich hyaline cartilage | ECM-rich hyaline cartilage | Muco-cartilage | Notochord | Fin Mes. | Sclero-tome | Pro-nephros | Somatic | Cardiac | Phar. endoderm | Oral ectoderm | CNS neurons | PNS neurons |
| **Gnathostomes** | *acan*<br>*vcan* | *acan*<br>*vcan* | N/A | *acan*<br>*bcan* | *acan* | *acan*<br>*vcan* | *vcan* | *acan*<br>*vcan* | *vcan* | *vcan* | *bcan* | *bcan*<br>*ncan* | *bcan*<br>*ncan*<br>*vcan* |
| **Lamprey** | *lecC*<br>*lecA* | ? | *lecA*<br>*lecB*<br>*lecD* | *lecA* | *lecA* | *lecA*<br>*lecB* | *lecB* | No *lec* | *lecA*<br>*lecD* | *lecB* | *lecB* | *lecA*<br>*lecB* | *lecA*<br>*lecB* |
| **Urochordate** | N/A | N/A | N/A | Present | N/A | N/A | N/A | N/A | Present | Present | Present | Present | Present |
| **Cephalochordate** | Present | N/A | N/A | Present | Present | Present | ? | Present | N/A | Present | Present | Present | Present |
| **Cell/tissue is unique to vertebrates?** | No | Yes | Yes | No | No | No | Yes<sup>4</sup> | No | No | No | No | No | No |
| **Cell/tissue has distinct histology in vertebrates?** | Yes<sup>1</sup> | - | - | No | Yes<sup>2</sup> | Yes<sup>3</sup> | - | Yes<sup>5</sup> | No | No | No | No | Yes<sup>6</sup> |

1. Unlike amphioxus cell-rich hyaline cartilage, vertebrate cell-rich hyaline cartilage develops from mesenchymal cells and can differentiate into ECM-rich hyaline cartilage
2. Amphioxus fin box mesoderm is epithelial and forms connective tissue, while vertebrate fin mesoderm delaminates and migrates as mesenchyme and can form cartilage and bone.
3. Amphioxus sclerotome is epithelial and forms connective tissue, while vertebrate sclerotome cells delaminate and migrate as mesenchyme and form cartilage and bone
4. Amphioxus nephridia and the vertebrate kidney consist of different cell types with no clear evolutionary relationship. The vertebrate kidney forms from migratory mesenchymal cells.
5. The amphioxus homolog of somatic LPM is epithelial and forms connective tissue, while gnathostome LPM cells delaminate and migrate as mesenchyme and form cartilage and bone in the limbs.
6. The PNS neurons of invertebrate chordates migrate short distances as neuroblasts, vertebrate PNS neurons migrate as multipotent mesenchymal cells that aggregate to form ganglia.

These comparisons reveal that *lectican* transcription is largely restricted to cells and tissues that are either *bona fide* vertebrate novelties, or have unique histology in vertebrates.

If *lectican*s are indeed expressed mainly in vertebrate histological innovations, what specifically did the first *lectican* contribute to the vertebrate phenotype? Perhaps the best studied Lectican is Aggrecan, which is a major component of the hyaline cartilage ECM, and confers many of its defining histological and structural properties. It was previously thought that hyaline cartilage was unique to vertebrates, though a clear homolog with virtually all of its defining features has recently been described in the invertebrate chordate amphioxus[99]. Thus, the origin of the first *lectican* was likely not prerequisite for the evolution of vertebrate-type cellular cartilage. Nevertheless, it is possible that *lectican*s contributed to the evolution of a more rigid type of cell rich hyaline cartilage, or the evolution of ECM-rich hyaline cartilage [Tab. 1].

Aside from hyaline cartilage, *lecticans* are expressed in a variety of other tissues during development. This suggests the evolution of the first *lectican* conferred a more general property upon vertebrate cells and tissues. Provocatively, a common theme among *lectican*-expressing cell types is that they differentiate from migratory and/or mesenchymal precursors [Tab. 1].

Furthermore, gnathostome *lectican*s have been shown to regulate the migration of neural crest cells, the major mesenchymal cell type in the nascent vertebrate head [28]. We thus speculate that the primordial Lectican may have functioned to promote mesenchymal histology and/or migratory behavior during development. In support of this scenario, development in invertebrate deuterostomes is largely, or completely epithelial. Indeed, the invertebrate with the most vertebrate-like body plan, amphioxus, develops without any discernible mesenchyme [100, 101]. Comprehensive analyses of *lectican* function in a wider range of vertebrates, including lamprey, using new methods for loss-of-function perturbation [102] should help test this hypothesis.

## ACKNOWLEDGEMENTS

The authors thank Scott Miehls at the USGS Hammond Bay Biological Station for providing adult sea lampreys. They also thank Jeremiah J. Smith at the University of Kentucky, Jr-Kai Yu at the Academia Sinica in Taipei, Taiwan, as well as Juan Pascual-Anaya and Shigeru Kuratani at the RIKEN Institute in Kobe, Japan for supplemental transcriptomic data. Zachary Root, Marek Romasek, Tyler Square, David Jandzik, and Daniel Medeiros were supported by National Science Foundation grants IOS 1656843 and IOS 1257040 to Daniel Medeiros. Zachary Root, Cara Allen, and Margaux Brewer were also supported by the Beverly Sears and EBIO grants through the University of Colorado Boulder. David Jandzik was additionally supported by the Scientific Grant Agency of the Slovak Republic VEGA grant No.1/0415/17.

## METHODS

## Isolation of lamprey *lectican* homologues

Lamprey *lectican* sequences were assembled from transcriptomic reads of Tahara t. 26.5 embryos and late larval oral disc tissue that were previously gathered. Sequences from these files were used for our phylogenetic and syntenic analyses. For *in situ* hybridizations for *lecA*, primers were designed from lamprey genomic sequence to amplify conserved exon sequences, which were cloned into the pJet1.2 vector from ThermoFisher©. For *lecB, lecC,* and *lecD*, 500bp regions from transcriptomic sequences were selected and ordered as fragments in pUC57-amp vector from Synbio Tech©.

### Phylogenetic Analysis

#### General Procedure

Peptide sequences of gnathostome genes were gathered on NCBI and aligned with lamprey genes using the PROBALIGN [71] program on CIPRES [73] servers. For all alignments, we used a gap open penalty of 20 and a gap extension penalty of 1. To determine the optimal substitution model for our phylogenetic analysis, we used ProtTest v3.4.2 [72]. For all tests, we allowed the possibility of invariant sites, empirical frequencies, and we used a fixed BIONJ tree topology to determine our ideal model. For our phylogenetic analysis, we used maximum likelihood analyses using RAxML-HPC2 Workflow on CIPRES servers. Using the parameters recommended by ProtTest, our likelihood scores were bootstrapped with 1000 trees for each test to derive a consensus tree. Our consensus trees were lastly visualized using FigTree v1. 4.4 [74].

#### Lectican Orthology Tests

Due to the large amounts of evolutionary time passing since the divergence of lamprey and gnathostomes, we tested both large and small numbers of taxa per gene as well as the inclusion or exclusion of hagfish sequences using the aforementioned methods. The *lectican* N terminus shares sequence similarities with the *hapln* genes, so we next performed phylogenetic analyses using the N termini of these genes, vertebrate *hapln* genes, as well as X-Link-containing genes that have been identified outside of vertebrates. Lecticans have overall more sequence conservation at the N and C termini, so we lastly tested substitution models that were specific to each termini, additionally removing the intermediate domain for these phylogenetic analyses. For our original *lectican* test, a DCMut + I + G + F model was calculated with a log likelihood score of -80,905.555 under Akaike Information Criterion (AIC). For our HAPLN test, a JTT + G model was calculated with a log likelihood score of -29,806.183 under AIC. Our *lectican* + hagfish test yielded a VT + I + G + F model with a log likelihood score of -152,828.685 under AIC. Our *lectican* + hagfish test yielded a VT + I + G + F model with a log likelihood score of -152,828.685 under AIC. Our HAPLN + N terminus test yielded a WAG + G + F model with a log likelihood score of -37,630.993 under AIC. When testing specific domain models, a WAG + I + G + F model was calculated for the N terminus with a log likelihood score of -37,763.00 under AIC. Conversely, our C terminus yielded a JTT + I + G model with a log likelihood score of -- 21.947.234 under AIC. Lastly, for our *mef2* test, a JTT+ G + F model was calculated with a log likelihood score of -18,094.435 under AIC.

#### Synteny Analysis

For our microsynteny analysis, we gathered peptide sequences of gnathostome *lectican*s on NCBI and found their respective genomic location using UCSC’s Genome Browser and BLAT tool [103, 104] as well as ENSEMBL [105]. We then used sequences of elephant shark (*Callorhinchus milii*), dog (*Canis lupus familiaris*), chicken (*Gallus gallus*), mouse (*Mus musculus*), African clawed frog (*Xenopus tropicalis*), Spotted gar (*Lepisosteus ocullatus*), and human (*Homo sapiens*) to reconstruct the ancestral arrangement of genes around the *acan, bcan, ncan,* and *vcan* loci. To do this, we compared genes in the +/- 250-300kb around each gnathostome locus. Syntelogs conserved in six of the seven gnathostomes (or five of seven gnathostomes, if one of the five organisms was elephant shark, the most basally diverging gnathostome analyzed), were included in the reconstructed loci. The orientation of each syntelog was determined by the majority orientation. We then used UCSC’s Genome Browser to identify all genes within +/- 400kb of each lamprey *lectican*. Because comparisons of the genes immediately adjacent to the gnathostome and lamprey *lectican* loci revealed very few conserved syntelogs [Fig. 2A], we expanded our analysis to include a larger selection of syntenic genes [Fig. S8]. To do this we identified the 20 genes (when available) immediately 5’ and 3’ of the *acan, ncan,* and *vcan* loci of three distantly related gnathostomes; chicken, elephant shark, and spotted gar. For *bcan*, which is not present in the current elephant shark genome assembly, we identified the 20 genes immediately 5’ and 3’ of the chicken, spotted gar, and zebrafish *bcan* loci. We then identified the 20 genes immediately 5’ and 3’ of lamprey *lecB and lecD*. For *lecC*, which sits near the 5’ end of a scaffold, we identified the seven 5’ genes, and the 33 3’ genes.

For *lecD,* which sits on a scaffold with less than 40 genes, all 29 flanking genes on the scaffold were used. For each gene identified as flanking a lamprey *lectican*, we then asked if there was a syntelog near one or more gnathostome *lecticans*. We then color-coded the lamprey genes based on the gnathostome *lectican(s)* its syntelog is linked to [Fig. S8]

### Embryo Collection and Staging

Embryos for in situ hybridization were obtained from adult spawning-phase sea lampreys (Petromyzon marinus) collected from Lake Huron, MI, and kept in chilled holding tanks as previously described [9]. Embryos were staged according to the method of Tahara [75], fixed in MEMFA (Mops buffer, EGTA, MgSO4, and formaldehyde), rinsed in Mops buffer, dehydrated into methanol, and stored at –20 °C.

### *In Situ* Hybridization

Riboprobes were made for anti-sense fragments using SP6 RNA Polymerase. Sequences for probes and genes are available upon request. In our experience, full-length P. marinus riboprobes, or riboprobes generated against untranslated regions of P. marinus transcripts, give higher background than short riboprobes against coding sequences. We believe that this is because lamprey noncoding sequences, especially 3′ UTRs, often have an excessive GC-repeat content, causing corresponding riboprobes to hybridize nonspecifically to off-targets. To mitigate this, we made short 500-bp riboprobes against coding regions and used a high-stringency hybridization protocol [95, 106]. Key parameters of this protocol include post-hybridization washes at 70 °C and the use of a low-salt, low-pH hybridization buffer (50% formamide; 1.3× SSC, pH 5.0; 5 mM EDTA, pH 8.0; 50 μg/mL tRNA; 0.2% Tween-20; 0.5% CHAPS; and 100 μg/mL heparin).

#### Histology and Sectioning

After in situ hybridization, embryos were postfixed in 4% paraformaldehyde/PBS (4 °C, overnight), rinsed in PBS, cryo-protected with 15% sucrose in water, embedded in 15% sucrose, 7.5% gelatin/15% sucrose (37 °C, several hours to overnight), and 20% gelatin/15% sucrose (37 °C overnight), frozen in -70 °C, and mounted with Tissue-Tek OCT compound (Sakura Finetek). Cryo-sections of 10 μm were collected on Super Frost Plus slides (Fisher Scientific), degelatinized in 3% gelatin in 38% ethanol, counterstained using Nuclear Fast Red (Vector Laboratories), dried, and cover-slipped with DPX (Fluka) [107].

#### Imaging

Whole-mount in situ hybridized *P. marinus* embryos and larvae were photographed using a Carl Zeiss Axiocam MRc5, Carl ZeissDiscovery V8 dissecting microscope, and Axiovision 4.6 software. Sections were photographed using a Carl Zeiss Imager A2 compound microscope.

---

**Figure S1.**
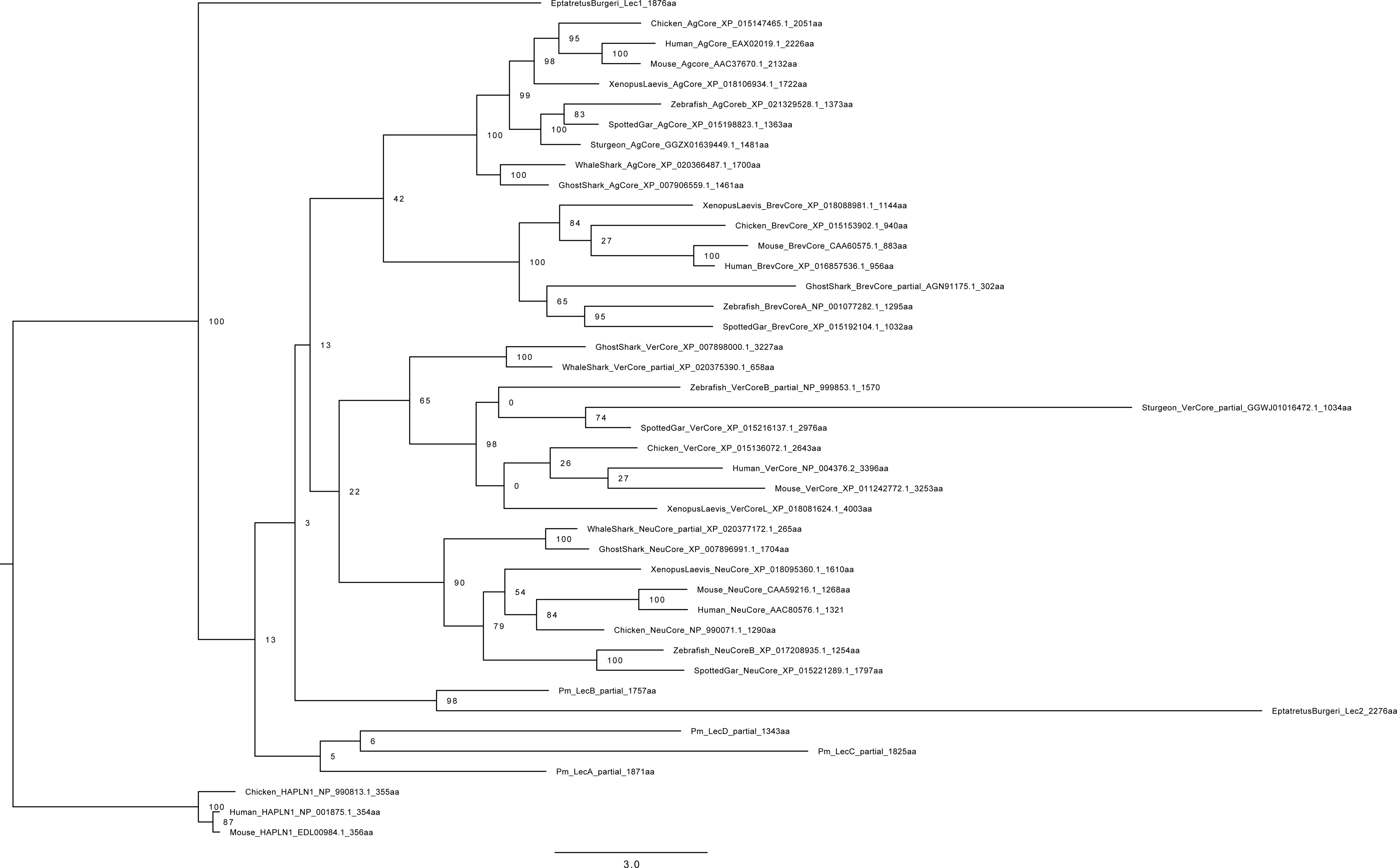
Phylogenetic tree built from lectican sequences in vertebrates with a larger number of taxa per gene as well as hagfish sequences. Maximum likelihood analysis scores are shown at the respective node. HAPLN1 sequences were designated as outgroup. Accession numbers for all sequences can be found in Tab. S1.

**Figure S2.**
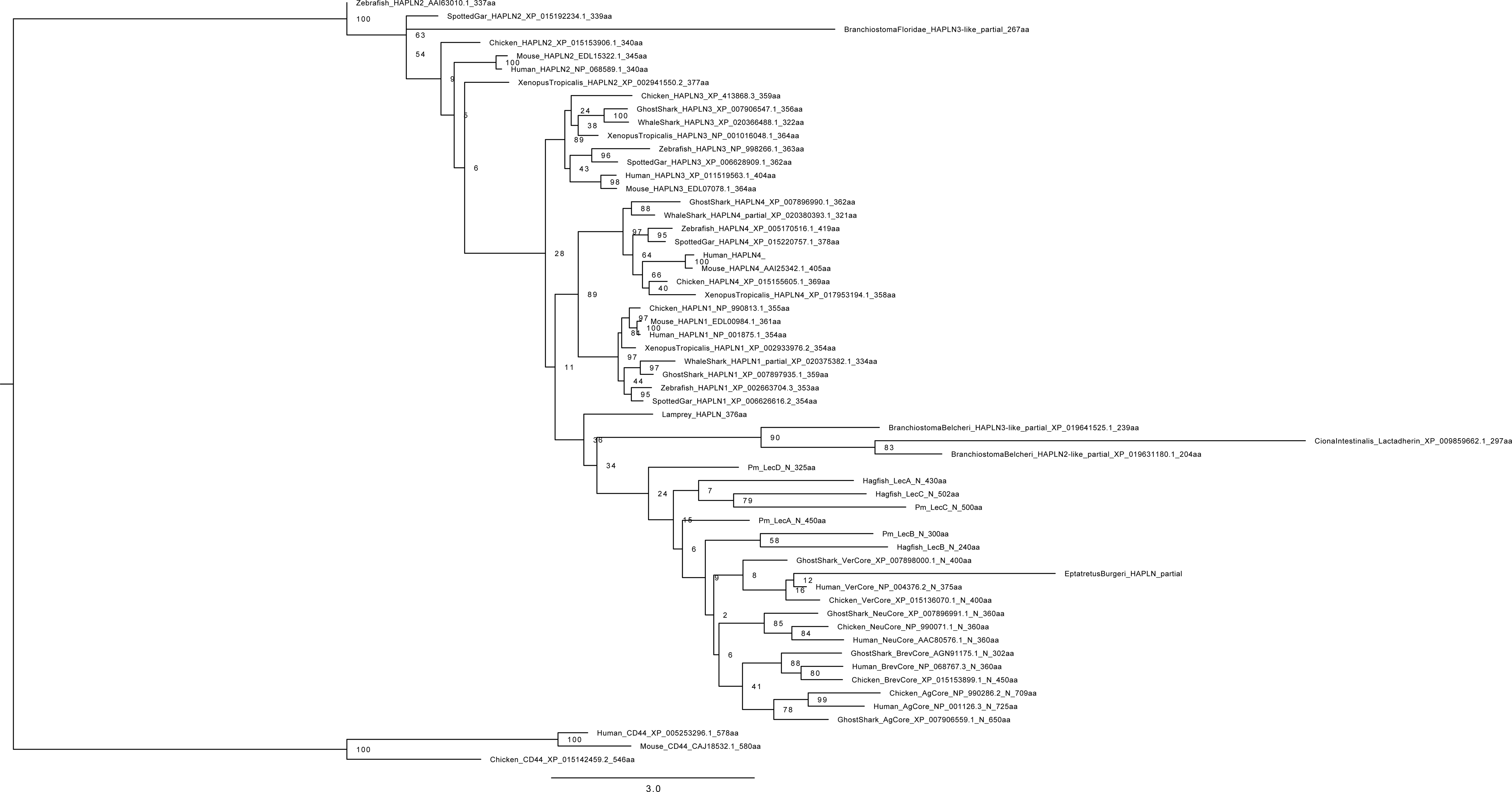
Phylogenetic tree built from lectican N-terminus sequences in vertebrates with a larger number of taxa per gene, hagfish N-terminus sequences, HAPLN genes, as well as X- link-containing genes outside of vertebrates. Maximum likelihood analysis scores are shown at the respective node. CD44 sequences were designated as outgroup. Accession numbers for all sequences can be found in Tab. S2.

**Figure S3.**
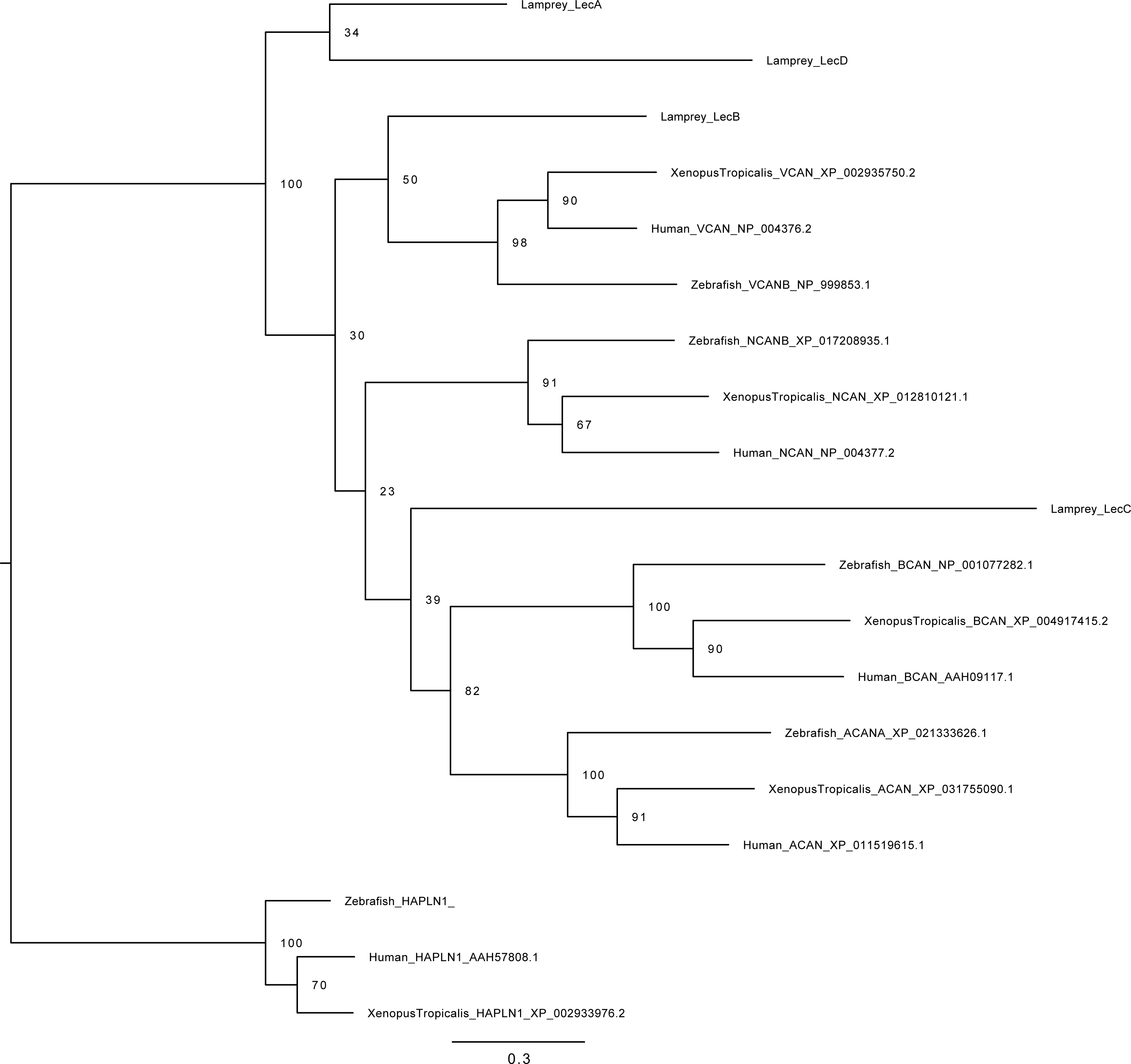
**P**hylogenetic tree built from lectican N+C termini sequences in gnathostomes and lamprey with minimal taxa and using the substitution model determined from the N-terminus. HAPLN1 sequences were designated as outgroup. Accession numbers for all sequences can be found in Tab. S3.

**Figure S4.**
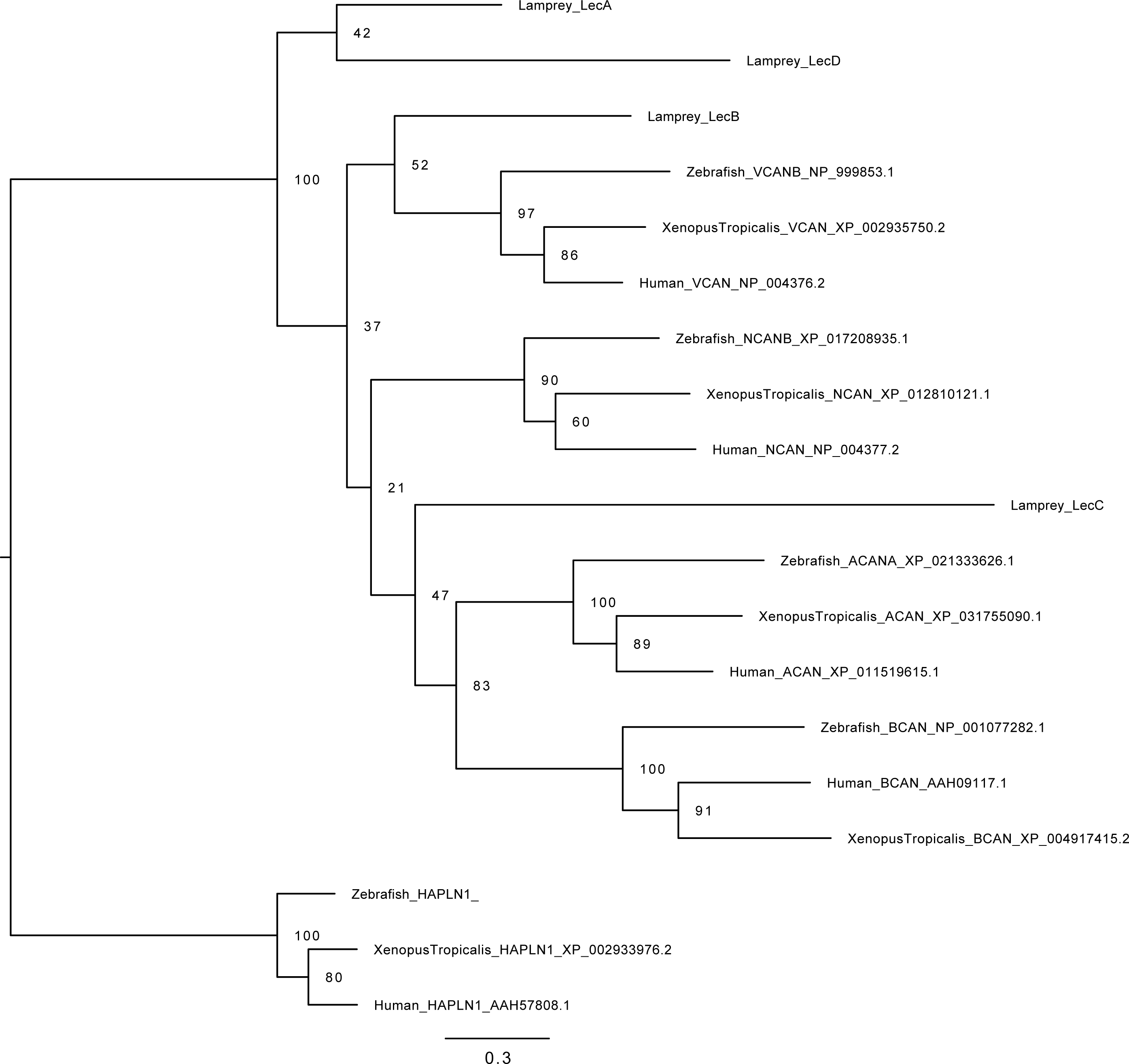
Phylogenetic tree built from lectican N+C termini sequences in gnathostomes and lamprey with minimal taxa and using the substitution model determined from the C-terminus. HAPLN1 sequences were designated as outgroup. Accession numbers for all sequences can be found in Tab. S3.

**Figure S5.**
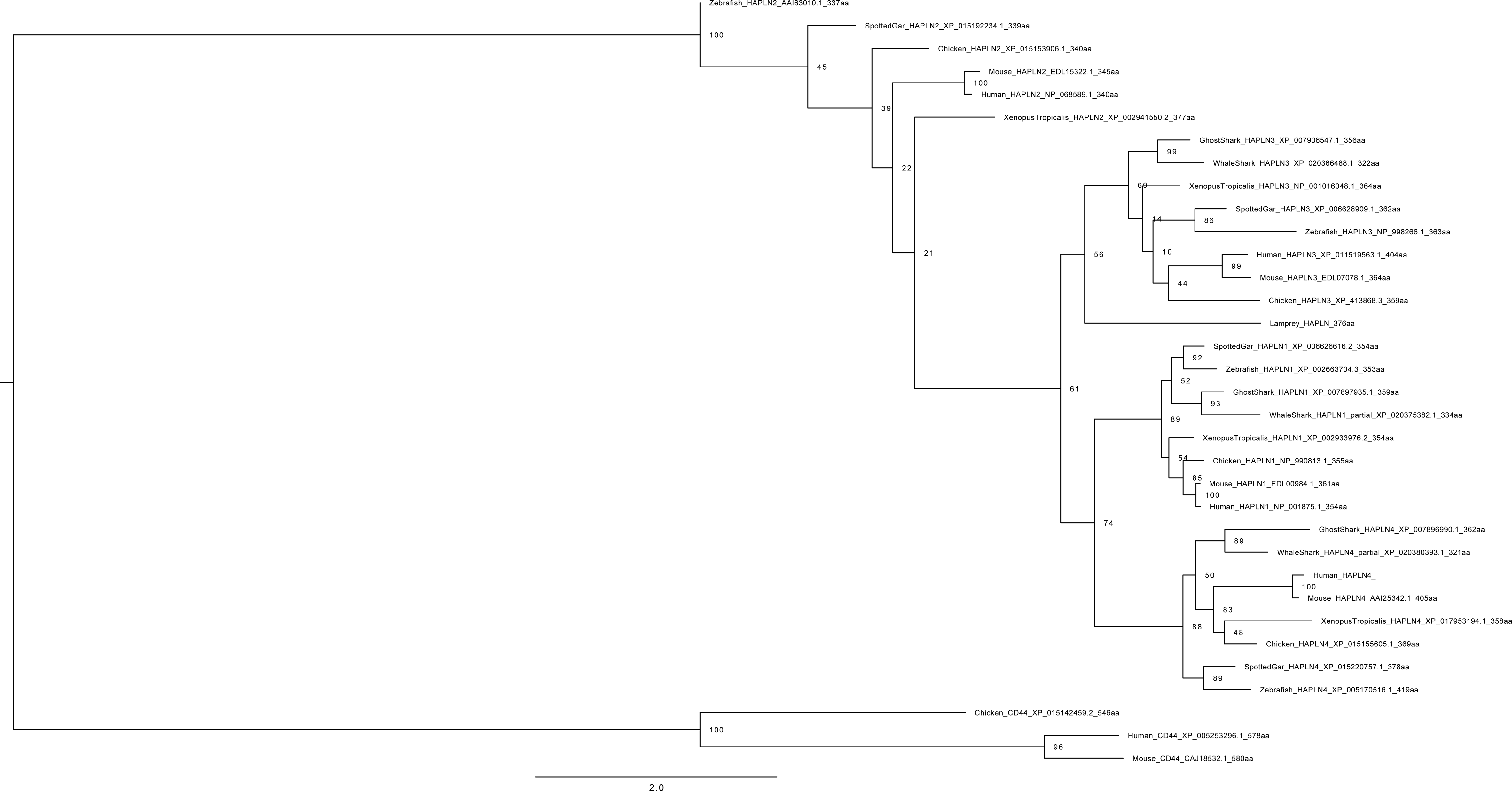
Final phylogenetic tree built from LECTICAN sequences in gnathostomes and lamprey. Maximum likelihood analysis scores are shown at the respective node. HAPLN1 sequences were designated as outgroup. Accession numbers for all sequences can be found in Tab. S4.

**Figure S6.**
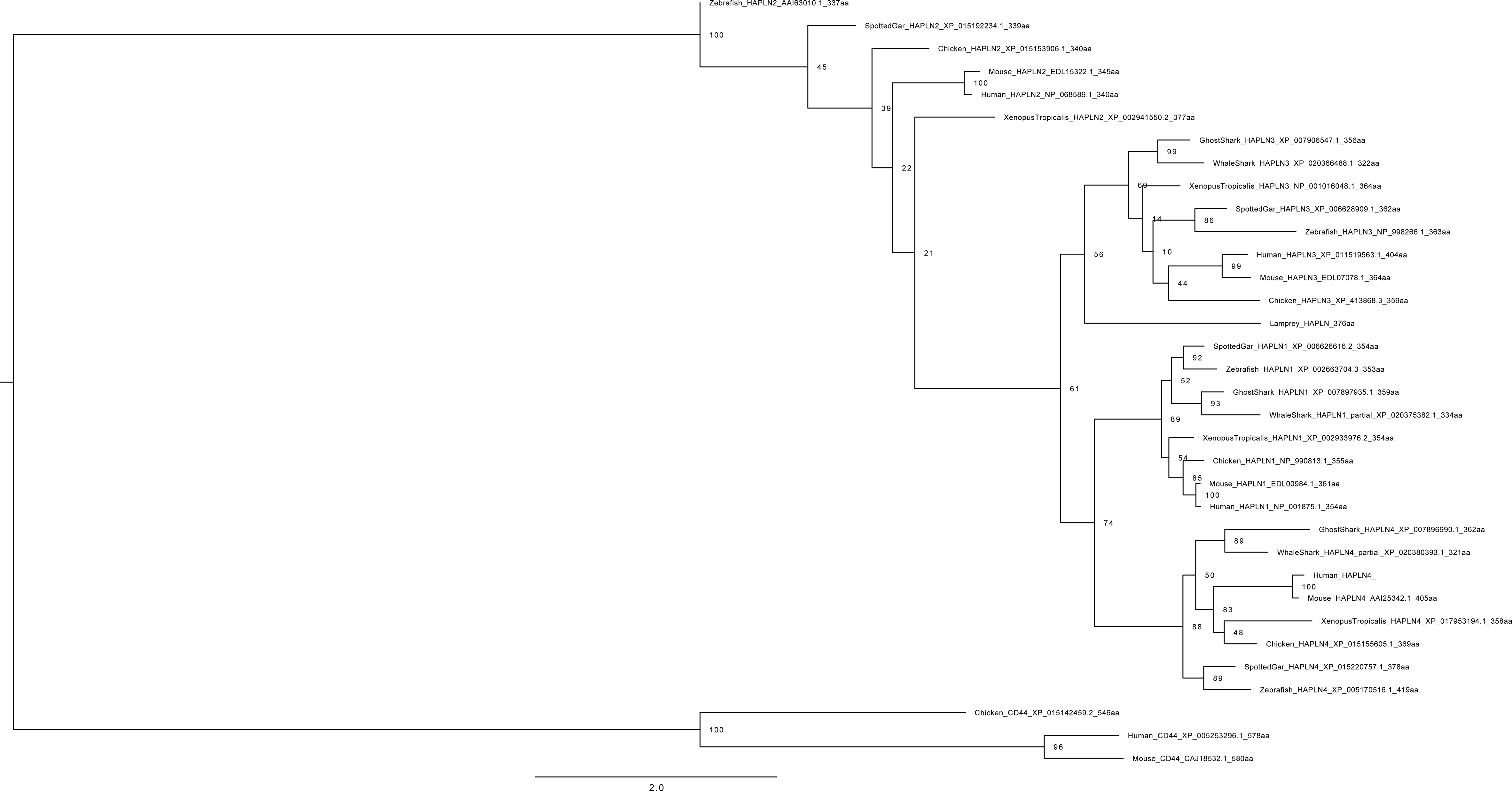
Phylogenetic tree reconstruction built from HAPLN sequences in gnathostomes and lamprey. Maximum likelihood analysis scores are shown at the respective node. CD44 sequences were designated as outgroup. Accession numbers for all sequences can be found in Tab. S5.

**Figure S7.**
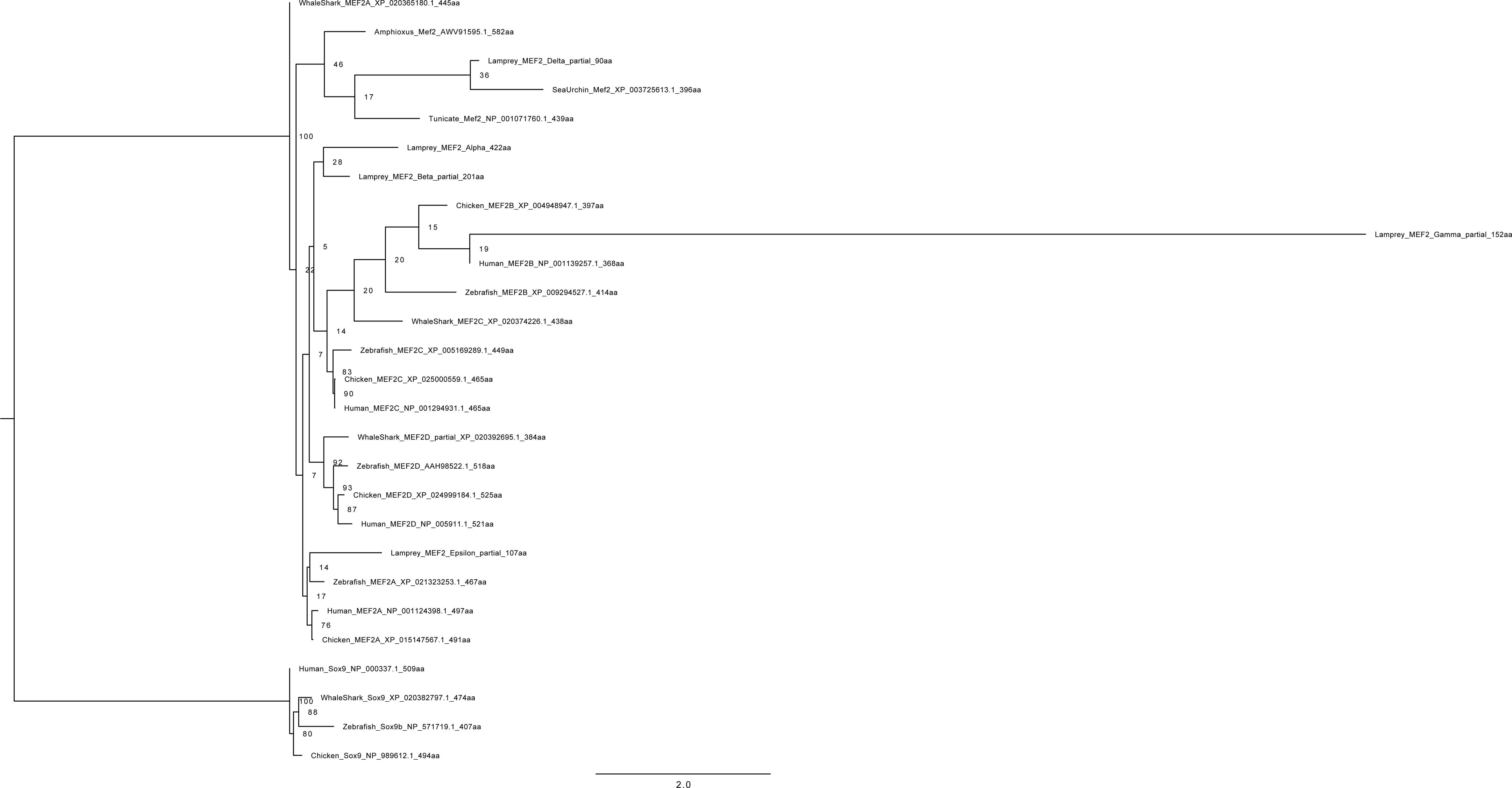
Phylogenetic tree reconstruction built from *MEF2* sequences in gnathostomes and lamprey. Maximum likelihood analysis scores are shown at the respective node. Sox9 sequences were designated as outgroup. Accession numbers for all sequences can be found in Tab. S6.

**Figure S8.**
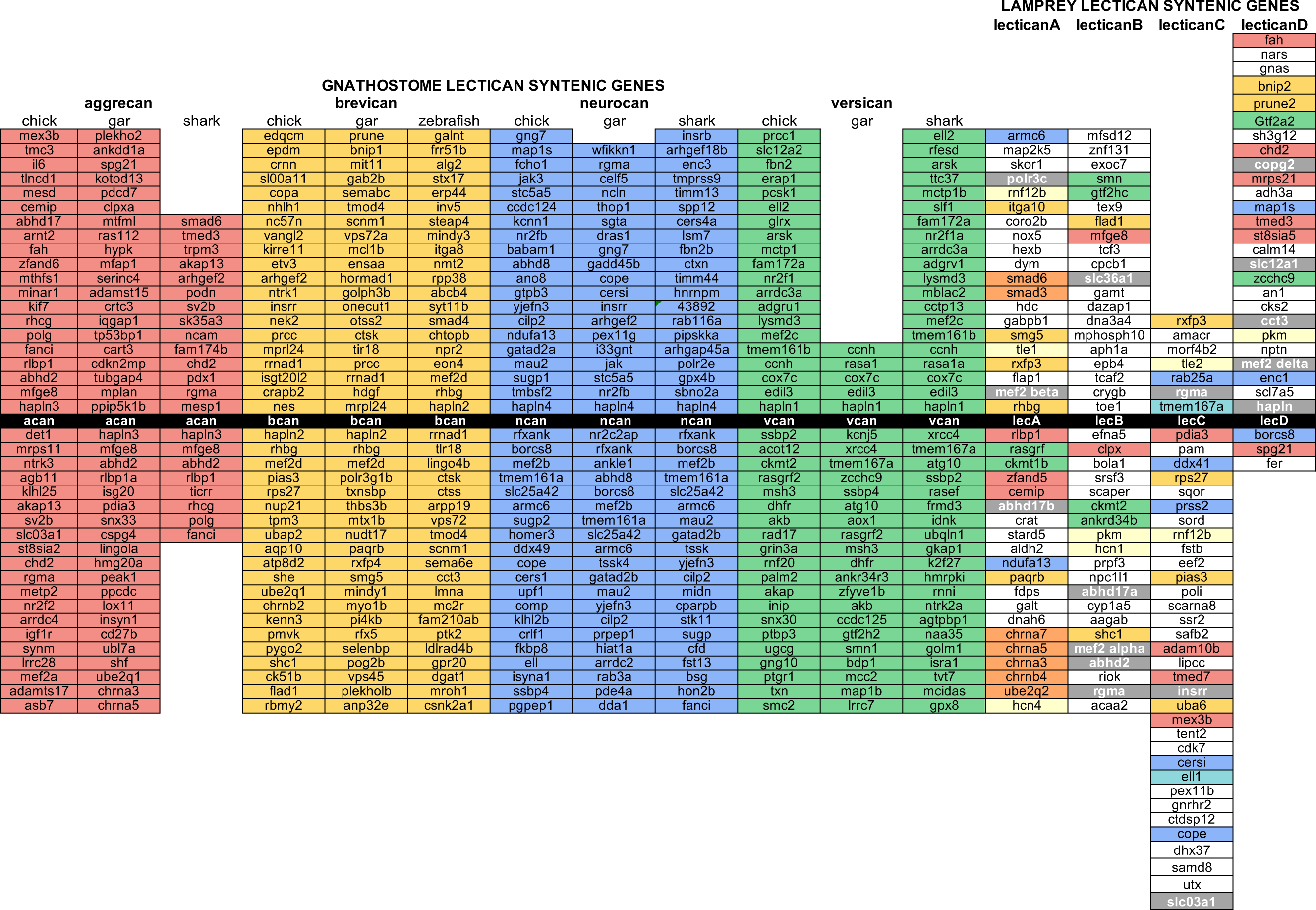
Comparison of genes syntenic to lamprey and gnathostome *lecticans*. Lamprey genes are color-coded based on the *lectican* their gnathostome syntelog is linked to. Red indicates the lamprey gene has a gnathostome homolog linked to *acan* only. Yellow indicates the lamprey gene has a gnathostome homolog linked to *bcan* only. Blue indicates the lamprey gene has a gnathostome homolog linked to *ncan* only. Green indicates the lamprey gene has a gnathostome homolog linked to *vcan* only. Orange indicates the lamprey gene has a gnathostome homolog linked to both *acan* and *bcan*. Turquoise indicates the lamprey gene has a gnathostome homolog linked to both *ncan* and *vcan*. Gray indicates the lamprey gene has gnathostome homologs linked to members of both the *acan+bcan* and *ncan+vcan* paralogy groups. Light yellow indicates lamprey genes with homologs linked to multiple lamprey *lecticans*. *LecA* is linked to 15 genes with gnathostome homologs linked exclusively to *acan* and/or *bcan*, but only 4 linked exclusively to *ncan* and/or *vcan*. The ratios of exclusive *acan* and/or *bcan* syntelogs to exclusive *ncan* and/or *vcan* syntelogs for *lecB*, *lecC*, and, *lecD* are 4:4, 8:7, and 8:5, respectively.

**Table S1.** NCBI accession numbers used for the phylogenetic analysis in Figure S1

| Sequence Name | Accession Number |
| --- | --- |
| Lamprey_HAPLN | MT125613 |
| Mouse_HAPLN1 | EDL00984.1 |
| ClawedFrog_HAPLN1 | XP_002933976.2 |
| GhostShark_HAPLN1 | XP_007897935.1 |
| WhaleShark_HAPLN1 | XP_020375382.1 |
| Human_HAPLN1 | NP_001875.1 |
| Chicken_HAPLN1 | NP_990813.1 |
| SpottedGar_HAPLN1 | XP_006626616.2 |
| Zebrafish_HAPLN1 | XP_002663704.3 |
| Mouse_HAPLN2 | EDL15322.1 |
| ClawedFrog_HAPLN2 | XP_002941550.2 |
| Human_HAPLN2 | NP_068589.1 |
| Chicken_HAPLN2 | XP_015153906.1 |
| SpottedGar_HAPLN2 | XP_015192234.1 |
| Zebrafish_HAPLN2 | AAI63010.1 |
| Mouse_HAPLN3 | EDL07078.1 |
| ClawedFrog_HAPLN3 | NP_001016048.1 |
| GhostShark_HAPLN3 | XP_007906547.1 |
| WhaleShark_HAPLN3 | XP_020366488.1 |
| Human_HAPLN3 | XP_011519563.1 |
| Chicken_HAPLN3 | XP_413868.3 |
| SpottedGar_HAPLN3 | XP_006628909.1 |
| Zebrafish_HAPLN3 | NP_998266.1 |
| Mouse_HAPLN4 | AAI25342.1 |
| ClawedFrog_HAPLN4 | XP_017953194.1 |
| GhostShark_HAPLN4 | XP_007896990.1 |
| WhaleShark_HAPLN4 | XP_020380393.1 |
| Human_HAPLN4 | NP_075378.1 |
| Chicken_HAPLN4 | XP_015155605.1 |
| SpottedGar_HAPLN4 | XP_015220757.1 |
| Zebrafish_HAPLN4 | XP_005170516.1 |
| Mouse_CD44 | CAJ18532.1 |
| Human_CD44 | XP_005253296.1 |
| Chicken_CD44 | XP_015142459.2 |

**Table S2.** NCBI accession numbers used for the phylogenetic analysis in Figure S2

|  |  |
| --- | --- |
| Lamprey_LecA | MT125609 |
| Lamprey_LecB | MT125610 |
| Lamprey_LecC | MT125611 |
| Lamprey_LecD | MT125612 |
| Lamprey_HAPLN | MT125613 |
| EptatretusBurgeri_Lec1 | MT559761 |
| EptatretusBurgeri_Lec2 | MT559762 |
| EptatretusBurgeri_Lec3 | MT582507 |
| EptatretusBurgeri_HAPLN | MT559763 |
| Amphioxus_HAPLN3-like_partial | MT582506 |
| Human_ACAN | EAX02019.1 |
| Chicken_ACAN | XP_015147465.1 |
| GhostShark_ACAN | XP_007906559.1 |
| Human_VCAN | NP_004376.2 |
| Chicken_VCAN | XP_015136072.1 |
| GhostShark_VCAN | XP_007898000.1 |
| GhostShark_BCAN | AGN91175.1 |
| Chicken_BCAN | XP_015153902.1 |
| Human_BCAN | XP_016857536.1 |
| Human_NCAN | AAC80576.1 |
| Chicken_NCAN | NP_990071.1 |
| GhostShark_NCAN | XP_007896991.1 |
| Mouse_HAPLN1 | EDL00984.1 |
| ClawedFrog_HAPLN1 | XP_002933976.2 |
| GhostShark_HAPLN1 | XP_007897935.1 |
| WhaleShark_HAPLN1 | XP_020375382.1 |
| Human_HAPLN1 | NP_001875.1 |
| Chicken_HAPLN1 | NP_990813.1 |
| SpottedGar_HAPLN1 | XP_006626616.2 |
| Zebrafish_HAPLN1 | XP_002663704.3 |
| Mouse_HAPLN2 | EDL15322.1 |
| ClawedFrog_HAPLN2 | XP_002941550.2 |
| Human_HAPLN2 | NP_068589.1 |
| Chicken_HAPLN2 | XP_015153906.1 |
| SpottedGar_HAPLN2 | XP_015192234.1 |
| Zebrafish_HAPLN2 | AAI63010.1 |
| Mouse_HAPLN3 | EDL07078.1 |
| ClawedFrog_HAPLN3 | NP_001016048.1 |
| GhostShark_HAPLN3 | XP_007906547.1 |
| WhaleShark_HAPLN3 | XP_020366488.1 |
| Human_HAPLN3 | XP_011519563.1 |
| Chicken_HAPLN3 | XP_413868.3 |
| SpottedGar_HAPLN3 | XP_006628909.1 |
| Zebrafish_HAPLN3 | NP_998266.1 |
| Mouse_HAPL4 | AAI25342.1 |
| ClawedFrog_HAPLN4 | XP_017953194.1 |
| GhostShark_HAPLN4 | XP_007896990.1 |
| WhaleShark_HAPLN4 | XP_020380393.1 |
| Human_HAPLN4 | NP_075378.1 |
| Chicken_HAPLN4 | XP_015155605.1 |
| SpottedGar_HAPLN4 | XP_015220757.1 |
| Zebrafish_HAPLN4 | XP_005170516.1 |
| Mouse_CD44 | CAJ18532.1 |
| Human_CD44 | XP_005253296.1 |
| Chicken_CD44 | XP_015142459.2 |
| Tunicate_Lactadherin | XP_009859662.1 |
| Amphioxus_HAPLN2-like_partial | XP_019631180.1 |
| Amphioxus_HAPLN3-like_partial | XP_019641525.1 |

**Table S3.**
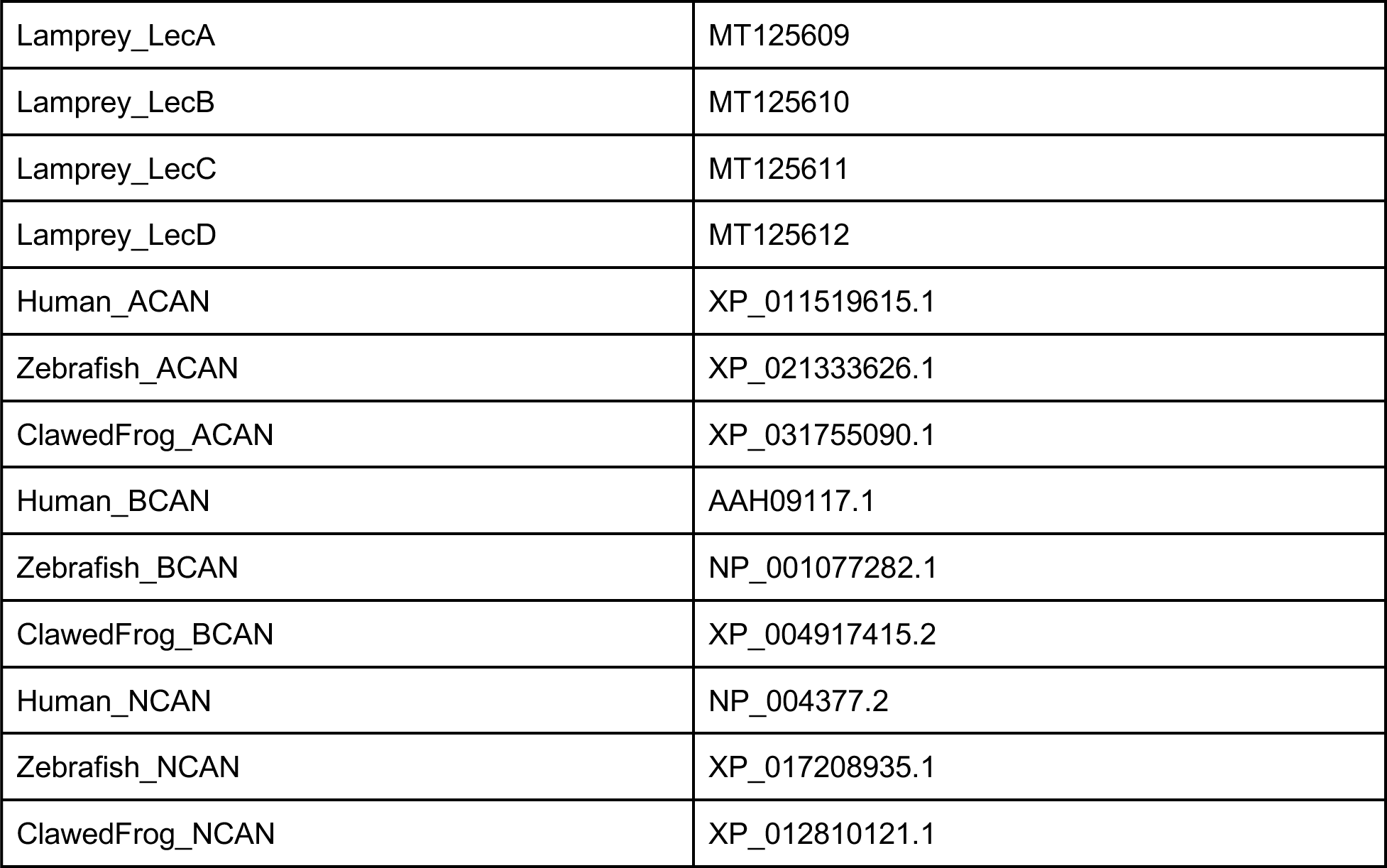
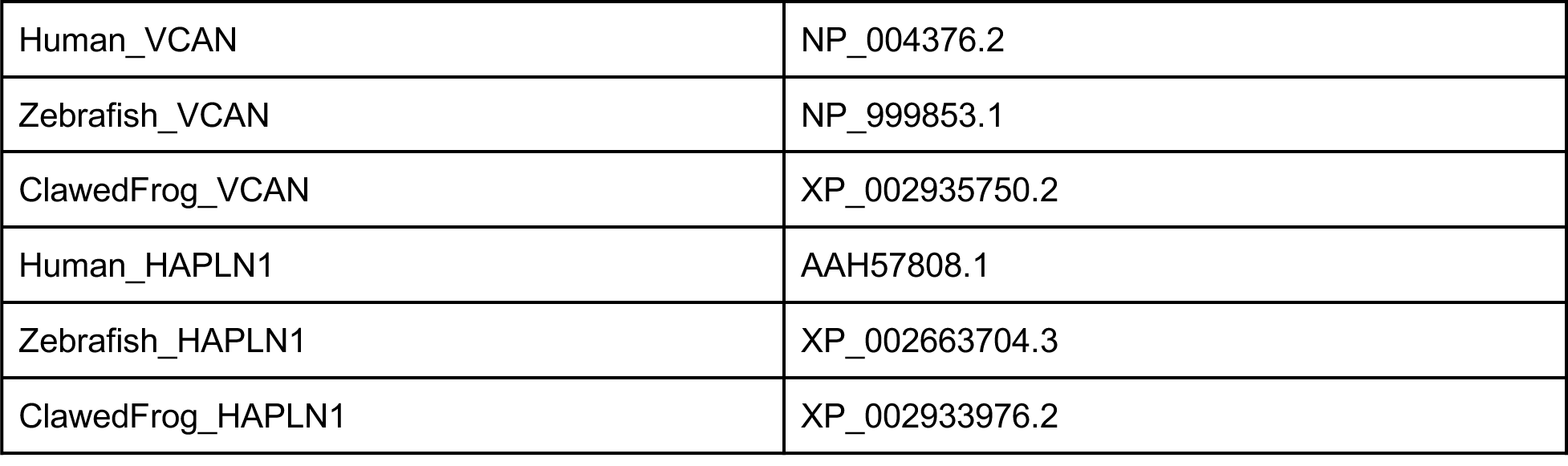
NCBI accession numbers used for the phylogenetic analysis in Figures S3 and S4

|  |  |
| --- | --- |
| Lamprey_LecA | MT125609 |
| Lamprey_LecB | MT125610 |
| Lamprey_LecC | MT125611 |
| Lamprey_LecD | MT125612 |
| Human_ACAN | XP_011519615.1 |
| Zebrafish_ACAN | XP_021333626.1 |
| ClawedFrog_ACAN | XP_031755090.1 |
| Human_BCAN | AAH09117.1 |
| Zebrafish_BCAN | NP_001077282.1 |
| ClawedFrog_BCAN | XP_004917415.2 |
| Human_NCAN | NP_004377.2 |
| Zebrafish_NCAN | XP_017208935.1 |
| ClawedFrog_NCAN | XP_012810121.1 |
| Human_VCAN | NP_004376.2 |
| Zebrafish_VCAN | NP_999853.1 |
| ClawedFrog_VCAN | XP_002935750.2 |
| Human_HAPLN1 | AAH57808.1 |
| Zebrafish_HAPLN1 | XP_002663704.3 |
| ClawedFrog_HAPLN1 | XP_002933976.2 |

**Table S4.** NCBI accession numbers used for the phylogenetic analysis in Figure S5

| Sequence Name | Accession Number |
| --- | --- |
| Lamprey_LecA | MT125609 |
| Lamprey_LecB | MT125610 |
| Lamprey_LecC | MT125611 |
| Lamprey_LecD | MT125612 |
| Human_ACAN | EAX02019.1 |
| Chicken_ACAN | XP_015147465.1 |
| Zebrafish_ACAN | XP_021329528.1 |
| GhostShark_ACAN | XP_007906559.1 |
| Human_VCAN | NP_004376.2 |
| Chicken_VCAN | XP_015136072.1 |
| Zebrafish_Dermacan | NP_999853.1 |
| GhostShark_VCAN | XP_007898000.1 |
| Zebrafish_VCAN | NP_001313486.1 |
| GhostShark_BCAN | AGN91175.1 |
| Chicken_BCAN | XP_015153902.1 |
| Human_BCAN | XP_016857536.1 |
| Zebrafish_BCAN | NP_001077282.1 |
| Human_NCAN | AAC80576.1 |
| Chicken_NCAN | NP_990071.1 |
| Zebrafish_NCAN | XP_017208935.1 |
| GhostShark_NCAN | XP_007896991.1 |
| Mouse_HAPLN1 | EDL00984.1 |
| Human_HAPLN1 | NP_001875.1 |
| Chicken_HAPLN1 | NP_990813.1 |

**Table S5.** NCBI accession numbers used for the phylogenetic analysis in Figure S6

|  |  |
| --- | --- |
| Lamprey_LecA | MT125609 |
| Lamprey_LecB | MT125610 |
| Lamprey_LecC | MT125611 |
| Lamprey_LecD | MT125612 |
| EptatretusBurgeri_Lec1 | MT559761 |
| EptatretusBurgeri_Lec2 | MT559762 |
| Human_ACAN | EAX02019.1 |
| Chicken_ACAN | XP_015147465.1 |
| Zebrafish_ACAN | XP_021329528.1 |
| GhostShark_ACAN | XP_007906559.1 |
| Mouse_ACAN | AAC37670.1 |
| ClawedFrog_ACAN | XP_018106934.1 |
| WhaleShark_ACAN | XP_020366487.1 |
| SpottedGar_ACAN | XP_015198823.1 |
| Sturgeon_ACAN | GGZX01639449.1 |
| Human_VCAN | NP_004376.2 |
| Chicken_VCAN | XP_015136072.1 |
| GhostShark_VCAN | XP_007898000.1 |
| Zebrafish_VCAN | NP_999853.1 |
| Mouse_VCAN | XP_011242772.1 |
| SpottedGar_VCAN | XP_015216137.1 |
| ClawedFrog_VCAN | XP_018081624.1 |
| WhaleShark_VCAN | XP_020375390.1 |
| Sturgeon_VCAN | GGWJ01016472.1 |
| GhostShark_BCAN | AGN91175.1 |
| Chicken_BCAN | XP_015153902.1 |
| Human_BCAN | XP_016857536.1 |
| Zebrafish_BCAN | NP_001077282.1 |
| Mouse_BCAN | CAA60575.1 |
| SpottedGar_BCAN | XP_015192104.1 |
| ClawedFrog_BCAN | XP_018088981.1 |
| Human_NCAN | AAC80576.1 |
| Chicken_NCAN | NP_990071.1 |
| Zebrafish_NCAN | XP_017208935.1 |
| GhostShark_NCAN | XP_007896991.1 |
| Mouse_NCAN | CAA59216.1 |
| ClawedFrog_NCAN | XP_018095360.1 |
| SpottedGar_NCAN | XP_015221289.1 |
| WhaleShark_NCAN | XP_020377172.1 |
| Mouse_HAPLN1 | EDL00984.1 |
| Human_HAPLN1 | NP_001875.1 |
| Chicken_HAPLN1 | NP_990813.1 |

**Table S6.** NCBI accession numbers used for the phylogenetic analysis in Figure S7

|  |  |
| --- | --- |
| Lamprey_Mef2Alpha | MT559756 |
| Lamprey_Mef2Beta | MT559757 |
| Lamprey_Mef2Gamma | MT559758 |
| Lamprey_Mef2Delta | MT559759 |
| Lamprey_Mef2Epsilon | MT559760 |
| Human_Mef2A | NP_001124398.1 |
| Chicken_Mef2A | XP_015147567.1 |
| Zebrafish_Mef2A | XP_021323253.1 |
| WhaleShark_Mef2A | XP_020365180.1 |
| Human_Mef2B | NP_001139257.1 |
| Chicken_Mef2B | XP_004948947.1 |
| Zebrafish_Mef2B | XP_009294527.1 |
| Human_Mef2C | NP_001294931.1 |
| Chicken_Mef2C | XP_025000559.1 |
| Zebrafish_Mef2C | XP_005169289.1 |
| WhaleShark_Mef2C | XP_020374226.1 |
| Human_Mef2D | NP_005911.1 |
| Chicken_Mef2D | XP_024999184.1 |
| Zebrafish_Mef2D | AAH98522.1 |
| WhaleShark_Mef2D | XP_020392695.1 |
| Tunicate_Mef2 | NP_001071760.1 |
| Amphioxus_Mef2 | AWV91595.1 |
| SeaUrchin_Mef2 | XP_003725613.1 |
| Human_Sox9 | NP_000337.1 |
| Chicken_Sox9 | NP_989612.1 |
| Zebrafish_Sox9 | NP_571719.1 |
| WhaleShark_Sox9 | XP_020382797.1 |

